# Trait Association and Prediction Through Integrative K-mer Analysis

**DOI:** 10.1101/2021.11.17.468725

**Authors:** Cheng He, Jacob D. Washburn, Yangfan Hao, Zhiwu Zhang, Jinliang Yang, Sanzhen Liu

**Affiliations:** Department of Plant Pathology, Kansas State University, Manhattan, KS 66506-5502, USA; Plant Genetics Research Unit, USDA-ARS, Columbia, MO 65211, USA; Department of Crop and Soil Sciences, Washington State University, Pullman, WA 99164, USA; Department of Agronomy and Horticulture, University of Nebraska-Lincoln, Lincoln, NE 68583-0915, USA; Center for Plant Science Innovation, University of Nebraska-Lincoln, Lincoln, NE 68583, USA

**Keywords:** k-mer, GWAS, maize, oil, leaf angle, prediction

## Abstract

Genome-wide association study (GWAS) with single nucleotide polymorphisms (SNPs) has been widely used to explore genetic controls of phenotypic traits. Here we employed an GWAS approach using k-mers, short substrings from sequencing reads. Using maize cob and kernel color traits, we demonstrated that k-mer GWAS can effectively identify associated k-mers. Co-expression analysis of kernel color k-mers and pathway genes directly found k-mers from causal genes. Analyzing complex traits of kernel oil and leaf angle resulted in k-mers from both known and candidate genes. Evolution analysis revealed most k-mers positively correlated with kernel oil were strongly selected against in maize populations, while most k-mers for upright leaf angle were positively selected. In addition, phenotypic prediction of kernel oil, leaf angle, and flowering time using k-mer data showed at least a similarly high prediction accuracy to the standard SNP-based method. Collectively, our results demonstrated the bridging role of k-mers for data integration and functional gene discovery.

## INTRODUCTION

Understanding the genetic basis of traits is a fundamental objective of biological studies. Genome-wide association study (GWAS) utilizing linkage disequilibrium (LD) between markers, largely single nucleotide polymorphisms, and causal variants has proven to be a valuable approach for genetic analysis, leading to the identification of a wealth of trait-associated loci (Visscher *et al*. 2017). The findings through GWAS have facilitated genetic dissection of complex human traits and diseases, translation to therapy applications, as well as plant and animal gene discovery and breeding (Liu and Yan 2019; Tam *et al*. 2019; Tian *et al*. 2020). However, also due to LD, which leads to spurious association, the approach typically is not able to directly pinpoint causal variants (Balding 2006). Traditionally, variant-trait associations rely on genotyping data that are based on reference genomes, and, mostly, derived from alignments of sequencing data. Reference genomes are known for their absence of sequences of non-reference haplotypes, presence of polymorphisms with other genomes, and even incompleteness, which altogether affect discovery of variants in diverse populations. In addition, alignment-based variant discovery often results in inaccurate genotyping owing to misalignments, particularly in complex genomes like maize (Lu *et al*. 2015).

GWAS can use counts of substrings of length *k* from longer sequencing reads, k-mers, as genotyping data (Rahman *et al*. 2018; Voichek and Weigel 2020). The k-mer occurrence count (KOC) from whole genome sequencing (WGS) data marks a range of genomic variants, including nucleotide substitutions, insertions, deletions, and copy number variants (CNVs). The KOC data are reference- and alignment-independent, solving some issues associated with the regular GWAS approach. Rahman et al. developed the analysis methodology HAWK (Hitting Associations With K-mers) that used KOCs to perform case-control associations (Rahman *et al*. 2018). In Voichek et al. 2020, the KOC of each k-mer was converted to presence and absence in each individual so that regular GWAS methods could be applied due to the similar structure to SNP genotyping data (Voichek and Weigel 2020). The conversion, however, removes information on any quantitative genetic variation that may exist, such as copy number variation.

K-mers can also be used for genome characterization and quantitative comparison between genomes. We have previously utilized low-pass whole genome sequencing reads from maize lines to examine evolutionary changes in highly repetitive sequences (Liu *et al*. 2017). We established a method to evaluate the quality of genome assemblies by comparing k-mer profiles in genome assemblies and original sequencing reads (He *et al*. 2020). In other studies, k-mers have been applied to quantify genome size and genome features such as repetitiveness and heterozygosity, as well as to perform genome comparisons (Liu *et al*. 2013; Anvar *et al*. 2014; Simpson 2014; Vurture *et al*. 2017; Sun *et al*. 2018). The simplicity of k-mer data provides an advantage for the integration of multiple sources of sequence data, and could potentially be used for trait predictions. Machine learning and deep learning models are effective and popular methods for predictive modeling across many areas of science. One area of machine learning that has been particularly successful, and has clear transferability to genetics, is natural language processing (NLP). In NLP, letters, words, sentences, or even whole documents are analyzed to determine the underlying linguistic meaning, gauge the sentiment of the author, or simply group words with similar usage patterns together. One relatively simple, yet effective, NLP method is commonly referred to as “Bag-of-words”. Bag-of-words models vectorize text into forms that can be processed by traditional statistical methods. Recently, “bag-of-words’’ was adapted for use in predicting the methylation status of different regions of plant genomes, which was termed “Bag-of-kmers” (Mejía-Guerra and Buckler 2019). Neural network and deep neural network approaches have also been applied to both natural language processing and the prediction of molecular and external phenotypes (Mejía-Guerra and Buckler 2019; Washburn *et al*. 2019, 2020, 2021).

In this study, we performed k-mer GWAS using normalized KOCs from WGS data of a diverse maize (*Zea mays* subsp. *mays*) panel. We integrated the k-mer GWAS result with co-occurrence of k-mers within the panel as well as co-expression of k-mers across hundreds of RNA-Seq datasets from multiple tissues, developmental stages, and treatments, in an attempt to reduce spurious association for candidate gene identification. We analyzed both simple and complex traits and showed the power of k-mer GWAS. The integration of k-mer GWAS results with k-mer data from evolutionary sequencing data depicts scenaria of historic selection on genetic variants of traits. In addition, we developed several NLP and neural network methodologies using k-mer data, a reference- and alignment-free genome predictive strategy, for phenotypic prediction.

## RESULTS

### K-mer occurrence counts (KOCs) in reads from diverse maize lines

Whole genome sequencing data from the Maize 282 Association Panel (maize282) (Flint-Garcia *et al*. 2005) were used to directly determine KOCs of 25-mers, of which *k* equals to 25. Raw k-mers were filtered to remove k-mers that are rare or had low-level variations in counts among lines (**Fig. 1a**, Methods), resulting in a total of 1.1 billion non-redundant k-mers from 261 inbred lines. Most lines contain 0.35 to 0.55 billion non-redundant k-mers (**Fig. 1b**). KOCs were normalized based on the total counts of a set of conserved single-copy k-mers, which were present only once in a genome and conserved across all or most genomes (Liu *et al*. 2017). The average total normal KOCs (total KOCs) of conserved single-copy k-mers is 1,108, which indicates that, if a k-mer is present in all lines and only once in each line, the expected occurrence count from all lines is 1,108. From the distribution of total KOCs, the most abundant total KOCs is 272 that is much smaller than 1,108, which indicates that a large number of k-mers are not present in all genomes of these maize lines (**Fig. 1c**).

**Fig. 1.**
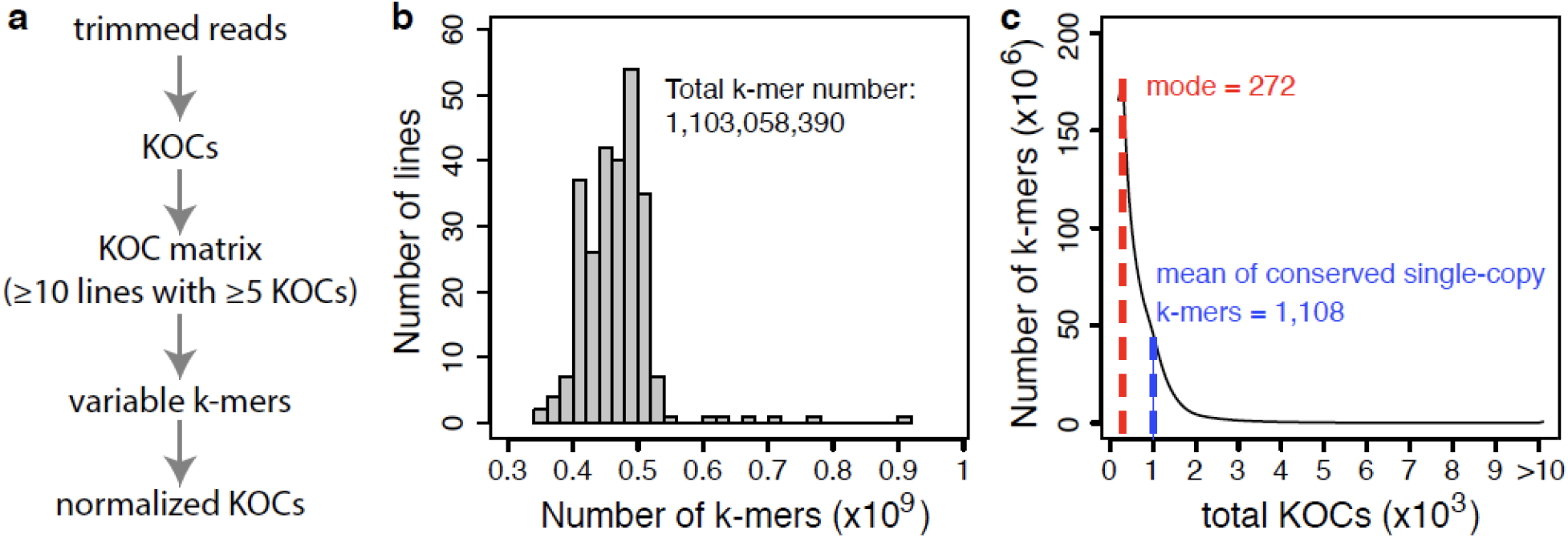
K-mer counts of maize inbred lines. (**a**) Flowchart for determining normalized k-mer occurrence counts (KOCs) for k-mer GWAS. (**b**) The distribution of k-mer numbers in 261 maize inbred lines. (**c**) The distribution of total KOC (total normalized k-mer occurrence count) per k-mer summed from 261 maize inbred lines.

### K-mer GWAS of cob color

Normalized KOCs from 1.1 billion k-mers were used as genotyping data to associate with the trait of cob color from a subset of maize282, of which 143 lines were categorized into: white (N=75) and red (N=68) (**Supplementary Fig. 1, Supplementary Table 1**). K-mer GWAS of cob color identified 16,383 associated k-mers with the p-value cutoff of 4.53e-11 (Bonferroni correction). To compare the result of SNP-based GWAS (SNP GWAS), k-mers were mapped to the B73 reference genome (Schnable *et al*. 2009; Jiao *et al*. 2017). Both GWAS results identified a strong association peak on the short arm of chromosome 1 (**Fig. 2a**, **2b**). *Pericarp color 1* (*P1*) encoding a MYB transcription factor located in the peak was known to regulate pigment genes (Grotewold *et al*. 1994). The B73 genome contains multiple copies of *P1* homologs. One of the *P1* homologs, Zm00001d028845, was hit by a set of k-mers with p-values smaller than 1e-22 (**Fig. 2c**). The average KOC of these k-mers, the mean depth of these k-mers, in red cob lines was approximately four times as high as that of the conserved single-copy k-mers, while the mean depth of these k-mers in white cob lines was close to zero (**Fig. 2d**). The result indicates that copy number variation of the *P1* gene is associated with cob color. Comparison of the haplotypes of Zm00001d028845 of red and white cob lines based on SNPs found that most red cob lines have the same haplotype, but the haplotypes of white cob lines are diverse and genotyping data of many SNPs are missing, which would reduce the statistical power of SNP GWAS (**Fig. 2e**). In addition to the Zm00001d028845 gene, another *P1* homologous gene, *Pericarp color 2* (*P2*), was also hit by associated k-mers but not by any markers in SNP GWAS (Zhang and Peterson 2005). Collectively, the cob color association analysis demonstrated the efficacy of k-mer GWAS and showed causal genes can be tagged by associated k-mers.

**Fig. 2.**
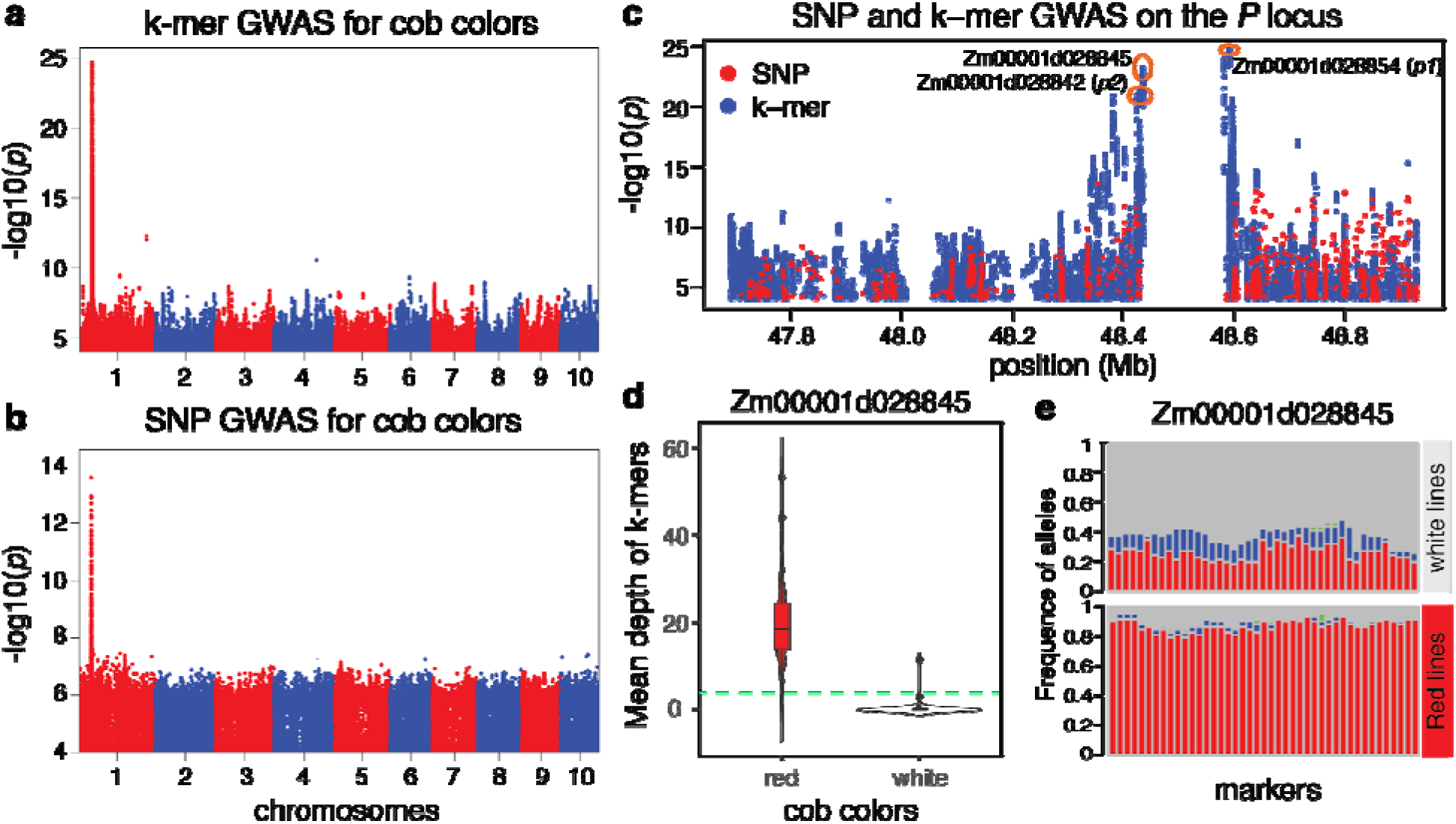
K-mer GWAS of cob color. (**a**) Manhattan plot of k-mer GWAS. (**b**) Manhattan plot of SNP GWAS. (**c**) Comparison of results from k-mer and SNP GWAS on the *p1*/*p2* locus. (**d**) Violin plot of mean depths, mean normalized k-mer occurrence counts (KOCs), of cob color k-mers from the gene Zm00001d028845 of lines with different cob colors. The green line indicates the average KOC of conserved single-copy k-mers. (**e**) Allele frequencies of SNPs on Zm00001d028845 of lines with different cob colors. Red, blue, green, and gray bars represent homozygotes of major alleles, homozygotes of minor alleles, heterozygotes, and missing data.

### K-mer GWAS of kernel colors

K-mer GWAS was performed using seed kernel colors from maize282, of which 259 lines were categorized into: white (N=66) and yellow (N=193) (**Supplementary Fig. 1**, **Supplementary Table 1**). The analysis resulted in 44,617 k-mers associated with kernel colors using the modified Bonferroni multiple test correction (p-value < 2.33e-08, Methods, **Data S1)**. To obtain genomic loci determining kernel color, trait-associated k-mers were clustered and assembled within a cluster. The resulting assembled contigs were then mapped to reference genomes. In detail, a co-occurrence network (CON) of kernel color associated k-mers was constructed based on their KOCs in 259 lines (Methods), which clustered k-mers with high correlations. This is similar to grouping SNP markers that are in high LD with each other. In total, 17 clusters including 58.8% of all associated k-mers were identified (**Data S1**). K-mers from each cluster were then assembled into contigs and aligned to the two reference genomes: the B73 reference genome and another maize inbred line A188 (Lin *et al*. 2021). As expected, most contigs in one cluster are co-located to a single locus on a chromosome. We defined such loci as the mapping locations of k-mer clusters. Most clusters were mapped to chromosomes 6 and 9 using either the B73 or A188 reference genomes, consistent with the result from SNP GWAS (**Figs. 3a**, **3b, 3c**). Two known kernel color genes, *yellow endosperm 1* (*y1*) (chromosome 6) and *carotenoid cleavage dioxygenase 1* (*ccd1*) (chromosome 9) were in these two loci (Buckner *et al*. 1990; Tan *et al*. 2017). Two other clusters were mapped to a region from 48.0 to 48.3 Mb on chromosome 2 of the A188 reference genome. The region covers a previously identified kernel color candidate gene, *zep1*, which encodes zeaxanthin epoxidase (Owens *et al*. 2014; Suwarno *et al*. 2015; Azmach *et al*. 2018). Another cluster was mapped on chromosome 3 but no candidate genes were identified. Mapping locations of three remaining clusters were not determined. Displaying correlations of randomly selected k-mers within clusters and among clusters supported the CON clustering results (**Fig. 3d**). Interestingly, relatively high correlations were observed among k-mers from chromosomes 2, 6, and 9, which harbor *zep1*, *y1*, and *ccd1*, respectively. Consistently, based on LD analysis, the SNPs from these three loci exhibited high correlations (**Supplementary Fig. 2**). Three top associated SNP sites on *zep1*, *y1*, and *ccd1* loci were used to examine allele combinations in the population. Approximately 69.7% (152/218) lines had the combination of the alleles corresponding to yellow color at all the three loci and produced yellow kernels, suggesting that these three loci were under parallel selection (**Fig. 3e**).

**Fig. 3.**
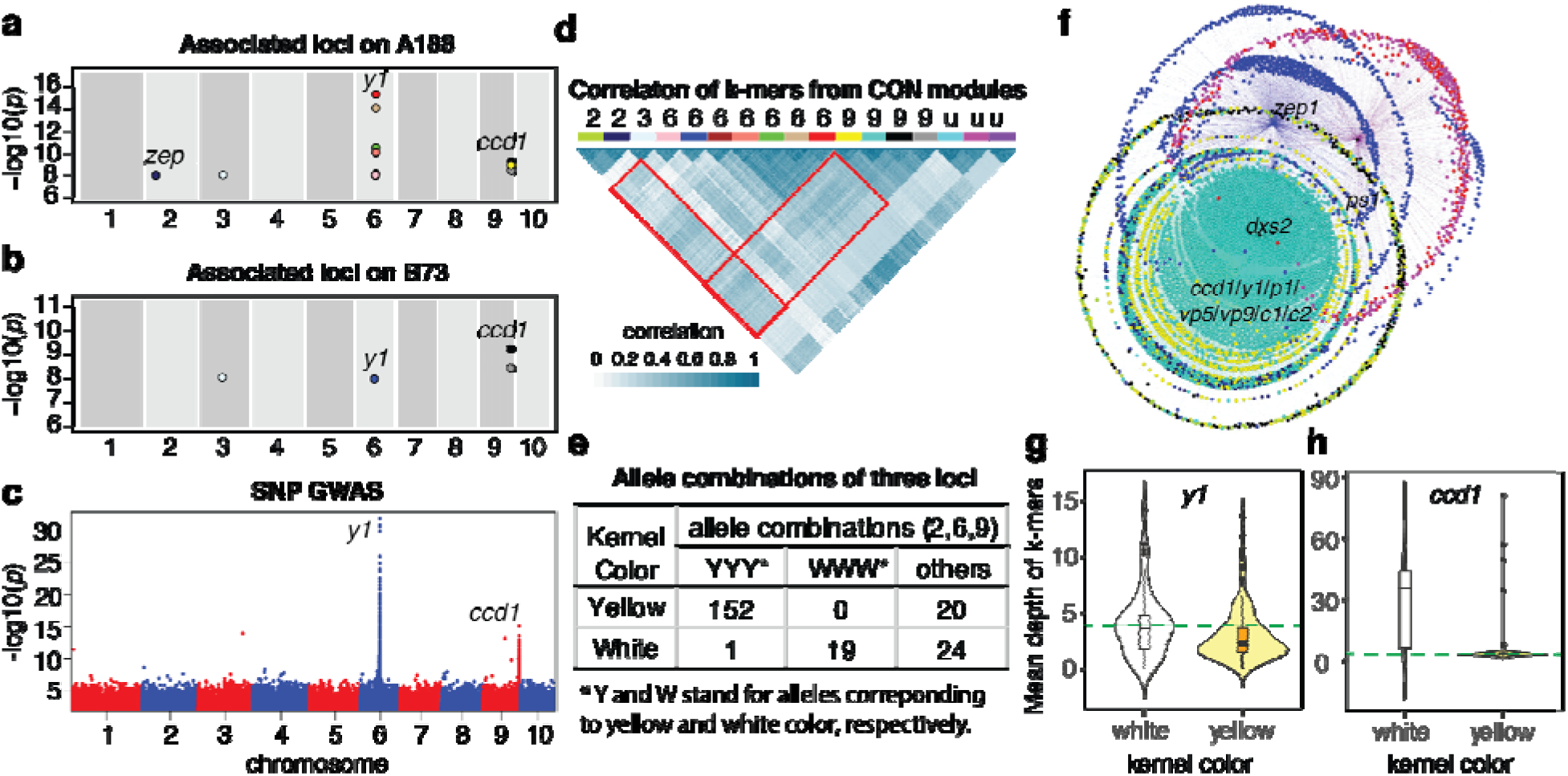
K-mer GWAS of kernel color. (**a**, **b**) Genomic locations of clusters of associated k-mers on the A188 (**a**) and B73 (**b**) reference genomes. Each point stands for a co-occurrence network (CON) cluster. Colors of points are consistent with cluster colors from CON analysis. (**c**) Manhattan plot of SNP GWAS. Two known kernel color genes, *y1* and *ccd1*, are marked around their genomic positions. (**d**) Correlation of 25 k-mers from each CON cluster based on normalized k-mer occurrence counts (KOCs). Colored bars represent CON clusters with consistent color codes. The mapped chromosome of each CON cluster is labeled. The letter “u” indicates that the cluster was not well mapped. (**e**) Three most associated SNP sites on the *zep1*, *y1*, and *ccd1* loci were analyzed. The alleles of each site were converted to Y and W that are corresponding to yellow and white color, respectively. The counts of allelic combinations were determined for lines with yellow seeds and lines with white seeds. Others stand for the combinations other than YYY or WWW. (**f**) Visualization of co-expression networks of kernel color k-mers overlapped with k-mers from RNA-Seq reads and known kernel color genes. Known genes in co-expression networks are labeled. (**g**, **h**) Violin plots of the mean depths, the mean normalized KOCs, of kernel color k-mers from the *y1* gene (**g**) and in the *ccd1* gene (**h**) of lines with different kernel colors. The green lines indicate the average KOC per line of conserved single-copy k-mers.

Kernel color associated k-mers were overlapped with k-mers derived from a large public RNA-Seq datasets (N=739, **Data S2**), all of which were generated using tissue samples from the reference inbred line B73. In total, 12,852 overlapping k-mers were identified. Expressions of these k-mers, the KOCs from RNA-Seq data, were used to build co-expression networks. In addition to these k-mers, 19 known seed color genes (**Supplementary Table 2**) are included in co-expression analysis. As a result, most k-mers co-expressed with known kernel color genes were clustered into 2 major modules turquoise and blue, including eight and two known genes, respectively (**Fig. 3f**). The result indicated that expressions of many seed color genes were co-modulated and the k-mers co-expressed with known genes are likely from genes conditioning seed color. The k-mers that are associated with kernel color and co-expressed with known seed color genes were then directly aligned to the B73 reference genome. Strikingly, 98.8% k-mers are located within *y1* and *ccd1*, both of which are among 19 known kernel color genes. We removed both *y1* and *ccd1* from the 19 known genes and repeated the co-expression analysis. Similarly, 99.7% of the resulting k-mers in networks are from *y1* and *ccd1*. The kernel color mapping results showed that integration of k-mer GWAS with co-expression network analysis could pinpoint causal genes in the cases that genes in the pathway are co-modulated to regulate phenotypic expression. Comparison of the mean depth of k-mers in the *y1* gene region between yellow and white kernel lines showed that both of them have a similar mean depth to conserved single copy k-mers (**Fig. 3g**). However, the mean depth of k-mers in the *ccd1* gene from white kernel lines were about 8 times higher than that of yellow kernel lines and 6 times higher than that of conserved single copy k-mers, implying that CNVs of the *ccd1* gene are likely associated with kernel color (**Fig. 3h**). This result is consistent with the previous findings that the regulative region or small polymorphisms of the *y1* gene influenced gene expression and condition kernel color (Lin *et al*. 2021), and the copy number of *ccd1* genes appeared to elevate the overall gene expression and whiten the kernel color (Tan *et al*. 2017; Lin *et al*. 2021).

### Optimization of the k-mer GWAS procedure

We compared 25-mer with 31-mer for the trait of kernel color. Using normalized KOCs, both analyses detected three loci: *y1*, *ccd1*, and *zep1* (**Supplementary Fig. 3**). Given that the longer mers have advantages in the assembly of k-mers into longer sequencers and direct mapping to the reference genomes, the 31-mer was used hereafter. We then compared four transformation methods using normalized KOCs (counts), log2 of counts, presence/absence of k-mers (presence/absence), and copy number of k-mers (copy number) as marker inputs for k-mer GWAS of kernel color. In the copy number method, the copy number of a k-mer represents the number of copies of each k-mer in the genome, which was inferred based on normalized KOCs. All four methods detected k-mers from the three known genes. The method of presence/absence, as compared to other methods, identified a slightly higher number of associated k-mers on *y1* and *zep1* whose causal variants do not belong to copy number variation (**Table 1**). However, for the *ccd1* gene whose copy number difference conditioning kernel color variation (Lin *et al*. 2021), the presence/absence method detected way less associated k-mers on the gene (**Table 1**). Out of the four methods, the copy number method exhibited a high power to detect associated k-mers from all three known loci. Therefore, the copy number method could serve as the method of selection or multiple methods could be combined, such as merging associated k-mers from both the presence/absence method and the copy number method to increase the GWAS power.

**Table 1.** Summary of kernel color k-mers from known genes from different methods

| Methods (31-mer) | Associated k-mers | *y1*<sup>1</sup> | *ccd1*<sup>1</sup> | *zep1*<sup>1</sup> |
| --- | --- | --- | --- | --- |
| counts | 51,526 | 427 | 5,509 | 54 |
| log2 of counts | 78,709 | 660 | 5,486 | 69 |
| presence/absence | 76,184 | 718 | 657 | 94 |
| copy number | 84,743 | 699 | 5,509 | 83 |

<sup>1</sup>at least 90% identity and >28 bp matches to *y1*, *ccd1*, *zep1* from B73v4

### K-mer GWAS of seed oil contents

Maize seeds with high oils are beneficial for food, feed and biofuel production (Li *et al*. 2013). The phenotype of kernel oil contents of most maize282 lines has been collected previously (**Supplementary Table 1)** (Peiffer *et al*. 2014). K-mer GWAS analysis of 259 maize282 lines identified 4,545 oil-associated k-mers (oil k-mers) with the p-value cutoff of 2.11e-8 (**Data S3**). Most oil k-mers (99.4%) are positively correlated with oil content, which were more mappable to the genome of the high-oil line P39 (41.9%, **Fig. 4a**) than to the genome of the low-oil line B73 (4.6%, **Fig. 4b**). From the P39 mapping result, peaks were identified on chromosomes 1, 2, 4, 6 and 7, which are supported by the quantitative trait loci (QTL) of oil data using Nested Association Mapping (NAM) Recombinant Inbred Line (RIL) populations (**Fig. 4c**, **Supplementary Table 3**).

**Fig. 4.**
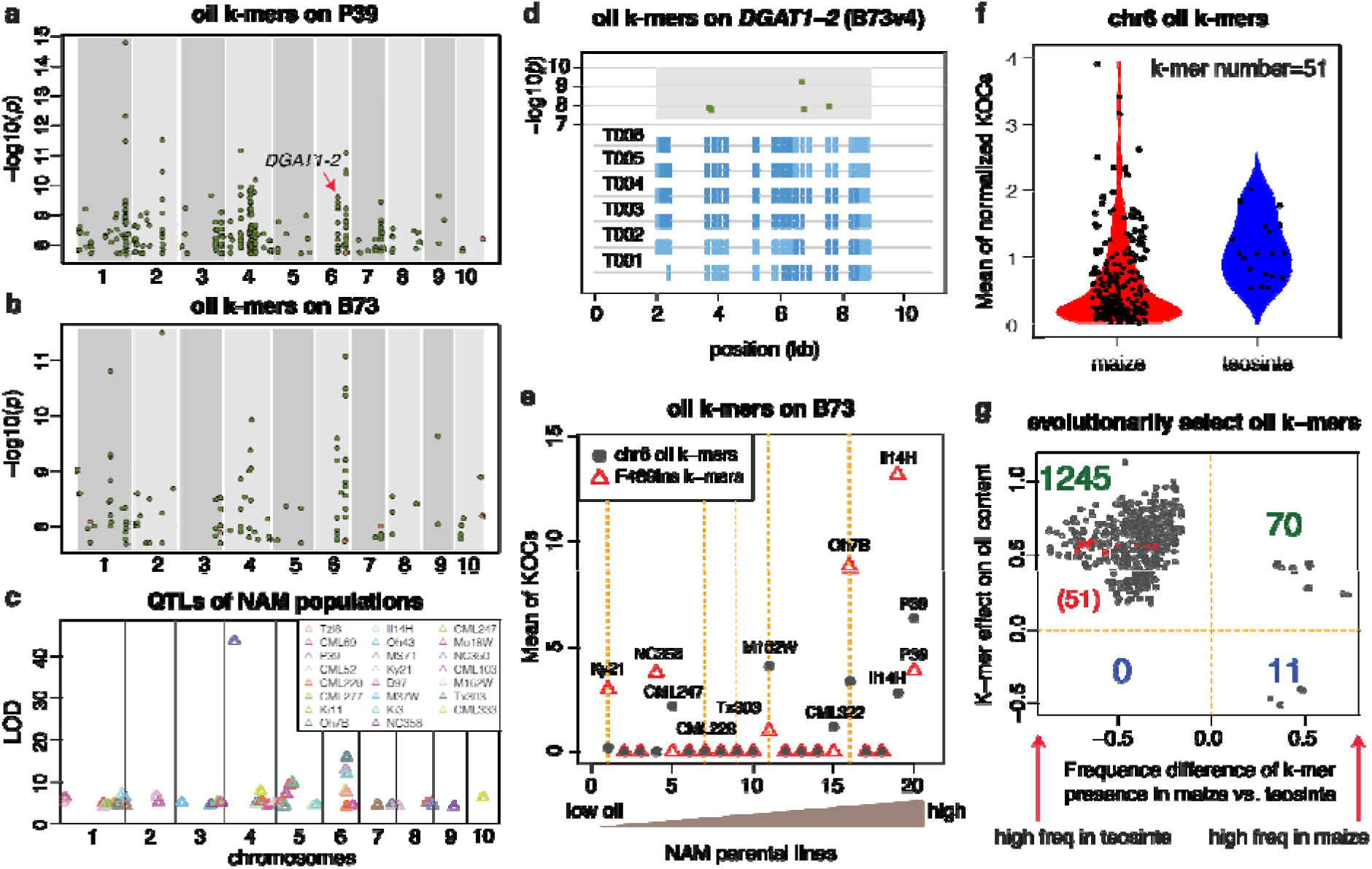
K-mer GWAS of kernel oil content. (**a**, **b**) Genomic locations of associated k-mers on the P39 (**a**) and B73 (**b**). Each point stands for an assembled k-mer contig or unassembled oil k-mers. Green and orange dots represent contigs from k-mers positively and negatively associated with kernel oil content, respectively. (**c**) QTL results of kernel oil in NAM populations. Each triangle represents the most significant marker for a QTL peak. (**d**) Genomic locations of oil k-mers on *DGAT1-2*. Dark blue boxes represent coding regions and light blue boxes represent untranslated regions (UTRs). Green squares stand for oil k-mers. (**e**) Normalized k-mer occurrence counts (KOCs) per NAM founder for the oil k-mers from the *DGAT1-2* locus on chromosome 6 and the k-mers from the F469ins region. (**f**) Comparison of average normalized KOCs of k-mers on *DGAT1-2* from lines of maize and teosinte. Each dot represents a maize or teosinte line. (**g**) Historically selected oil k-mers in maize and teosinte. Numbers of k-mers in each group are listed. K-mers from *DGAT1-2* are marked with red color.

The known gene *DGAT1-2* conditioning oil content is located on the chromosome 6 peak (Zheng *et al*. 2008). In the oil QTL analysis of NAM subpopulations, of which B73 is the common parent, the QTL of the *DGAT1-2* locus was identified in the subpopulations from six non-B73 parents, namely, M162W, Tx303, Tzi8, Oh7B, CML228 and Ky21. Comparison of the DGAT1-2 protein sequences with B73 showed all these lines carry a variant, F469ins, which has an amino acid insertion at the position 469 of the B73 protein sequence (**Supplementary Fig. 4a**). F469ins has been proved to confer a 27-37% increase in oil content (Zheng *et al*. 2008; Chai *et al*. 2012). Most k-mers derived from this causal polymorphic site, referred to as F469ins k-mers, had p-values ranging from 5.1e-5 to 8.5e-7 from k-mer GWAS but did not pass the significant threshold to be oil k-mers, which is likely due to some low-oil lines (e.g., Ky21 and NC358) containing F469ins (**Supplementary Fig. 4b**). Instead, k-mers from other polymorphic sites from the gene had significant p-values, which seem to be more predictive for oil contents than F469ins k-mers (**Fig. 4d**,**4e)**.

The kernel oil content is well known to be negatively correlated with the maize yield (Abou-Deif *et al*. 2012), indicating that the oil associated loci may be under negative selection. We compared the presence of the oil k-mers on chromosome 6 in lines of maize and teosinte (*Zea mays* subsp. *parviglumis*), the relative to the maize ancestor, using whole genome sequencing data of 257 maize282 lines and 20 diverse teosinte lines (Xu *et al*. 2020) (**Data S3**). The comparison clearly showed that the chromosome 6 oil k-mers are present in most teosinte lines but absent in most modern inbred lines (**Fig. 4f**). A similar pattern was observed for F469ins k-mers (**Supplementary Fig. 5**). The result is highly consistent with the previous report that the *DGAT1-2* allele with F469ins is ancestral, whereas the allele without F469ins was resulted from a more recent mutation selected during maize domestication and/or improvement (Zheng *et al*. 2008). We then extended the examination through a *χ*^2^ statistical test for each oil k-mer. The hypothesis is that the proportion of lines carrying a given k-mer in 20 teosinte lines is not different from that in 257 maize282 lines. As a result, 1,326 oil k-mers were identified as historically selected oil k-mers, each of which had differential representative proportions in teosinte and maize populations (FDR < 0.05, **Supplementary Table 4**). Strikingly, most historically selected oil k-mers (1,245/1,326) are present at a higher frequency in teosinte lines than in modern maize lines and almost all of them (1,315/1,326) are positively correlated with the oil content (**Fig. 4g**). The result supports that the high-oil trait was strongly selected against during maize domestication and/or improvement.

### K-mer GWAS of upper leaf angle

Maize leaf angle is a critical trait that may determine the optimization of planting density. A desirable plant architecture with an upright leaf angle of top leaves enhances the light capture efficiency for photosynthesis, thereby increasing maize grain yield (Pendleton *et al*. 1968; Duncan 1971). The trait of upper leaf angle (ULA) of the maize282 lines has been previously collected (Peiffer *et al*. 2014). K-mer GWAS using the ULA data from 259 lines identified 9,169 ULA associated k-mers (p-value < 2.82e-7) (**Data S4**). Mapping of associated k-mers, or ULA k-mers, to the B73 genome resulted in three strong peaks on chromosomes 3, 8, and 10 (**Supplementary Fig. 6**). We found the KOCs of ULA k-mers on chromosome 10 are highly correlated with the population structure. Therefore, we added the KOCs of the top ULA k-mer with the smallest p-value of chromosome 10 as a co-variant in k-mer GWAS. Reanalysis with the modified model resulted in additional ULA k-mers that can be mapped to the peaks on chromosomes 2, 3, 4, 5, 7, and 9 (**Fig. 5a**). Of these peaks, the peaks on chromosomes 3, 5 and 7 were supported by the ULA QTL analysis using NAM populations (**Supplementary Fig. 7, Supplementary Table 5**). In particular, the QTLs overlapping with the chromosome 3 peak were detected in more than half (11/19) NAM subpopulations and the underlying gene is unknown.

**Fig. 5.**
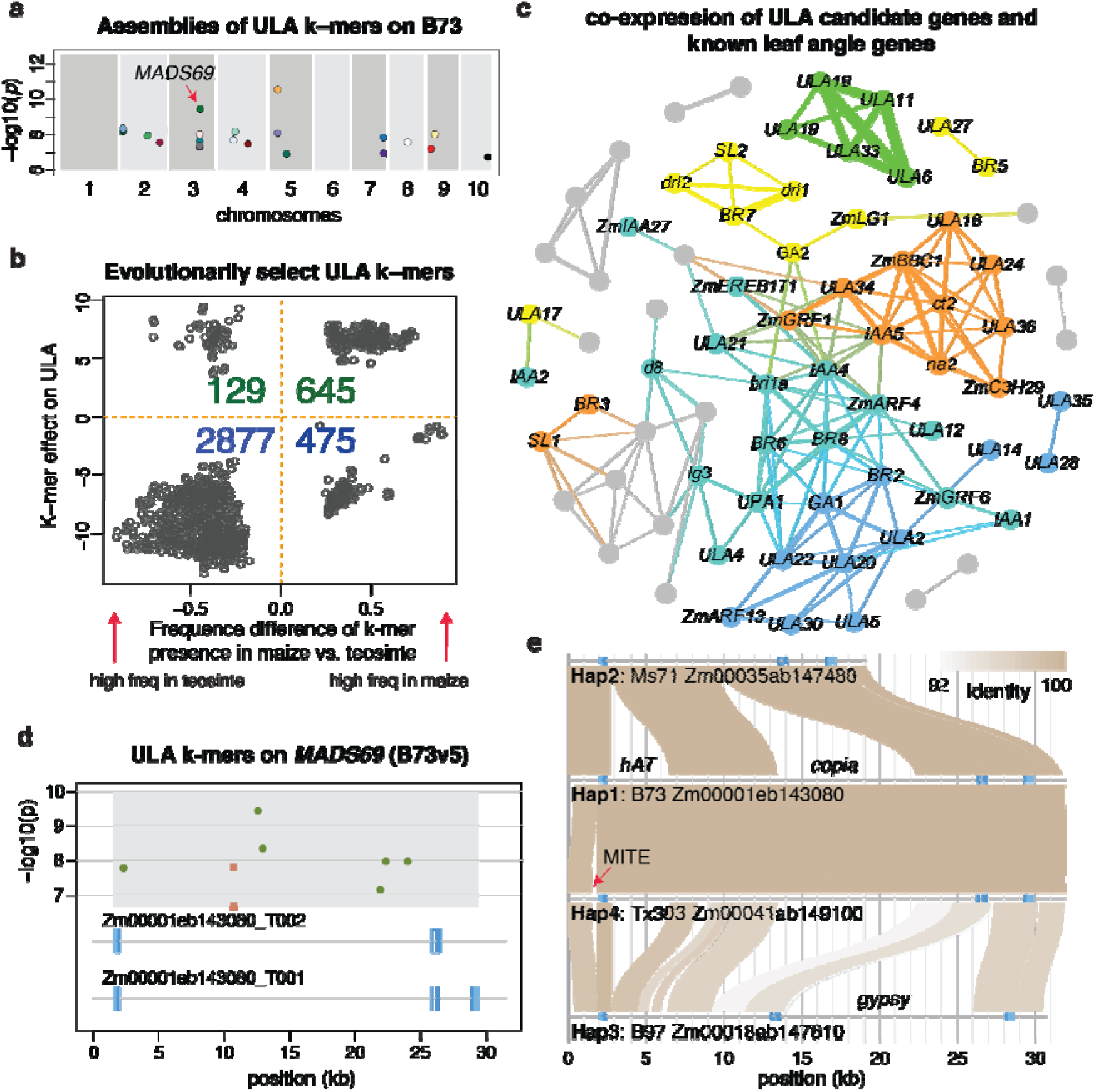
K-mer GWAS of upper leaf angle (ULA). **(a)** Locations of ULA k-mer clusters on the B73 reference genome. Each point stands for a cluster in the co-occurrence network (CON). **(b)** Historically selected ULA k-mers in maize and teosinte. **(c)** Co- expression network analysis of candidate genes and known ULA genes. Colors indicate different modules. **(d)** Location of ULA k-mers on *MADS69*. Dark blue boxes represent coding regions and light blue boxes represent UTRs. Green squares stand for the k- mers positively associated with upright ULA and orange squares stand for those negatively associated with upright ULA. (**e**) Four haplotypes (Hap1, 2, 3, 4) of *MADS69* from NAM parental lines. Annotated transposable elements are labeled.

Leaf angle is the trait that was under selection during maize breeding (Ariyanayagam *et al*. 1974; Pepper *et al*. 1977). We hypothesized that the genomic variants corresponding to the desirable and undesirable leaf angle are over- and under- represented in the population of modern maize lines, respectively. To test the hypothesis, the presence of ULA k-mers using WGS data of 20 teosinte lines (Xu *et al*. 2020) were compared to their presence in maize282 through the - χ statistical test. In total, we found 4,136 ULA k-mers are unproportionally presented in maize and teosinte populations, which are deemed to be historically selected ULA k-mers (**Fig. 5b, Supplementary Table 6**). Of the favorably selected ULA k-mers, which positively contribute to the upright leaf phenotype, 83.3% (645/774) are enriched in the maize population; while 85.6% of 3,362 unfavorably selected ULA k-mers are enriched in the teosinte population. The evolutionary data support that ULA k-mers were under strong selection. With mapping of selected ULA k-mers on the B73 reference genome (B73v4), 37 candidate genes were identified (**Supplementary Table 7**).

To prioritize these genes, co-expression analysis was performed between these 37 candidate genes and 55 leaf angle related genes using 739 B73 RNA-Seq data (**Fig 5c**, **Supplementary Table 8**) (Tian *et al*. 2019; Li *et al*. 2020). Note that genes instead of k-mers were used for co-expression analysis because most ULA k-mers did not match k-mers from B73 RNA-Seq reads. We found that 26 ULA candidate genes co-expressed with 48 leaf angle related genes. In the largest turquoise module, three ULA candidate genes, Zm00001d002425 (*ULA4*), Zm00001d042315 (*ULA12*) and Zm00001d051893 (*ULA21*), were co-expressed with 13 known ULA genes, including *liguleless3* (*lg3*) and *liguleless4* (*lg4*) (Muehlbauer *et al*. 1997). Most leaf angle related genes of the turquoise module are involved in phytohormone pathways, such as Auxin (*IAA, ZmARF4, ZmGRF1, ZmGRF6, ZmIAA27*), Brassinolide (*BR, bri1a*) and Gibberellin (*d8*) (van Zanten *et al*. 2010; Luo *et al*. 2016; Liu *et al*. 2019). ULA12 (Zm00001d042315) in the turquoise module encodes a transcription factor (TF) MADS69 that regulates flowering time of maize, and elevated expression levels of *MADS69* were shown to shorten maize flowering time (Liang *et al*. 2019). The *MADS69* gene was located in the QTL on chromosome 3 from more than half (11/19) NAM subpopulations (**Supplementary Fig. 7**). The ULA k-mers are located in the first intron of the B73 *MADS69* gene (**Fig. 5d**). Here, the version 5 B73 reference genome (B73v5) was used due to the incorrect annotation of this gene in B73v4. Using the recent genome assemblies of NAM founders and A188 (Lin *et al*. 2021; Hufford *et al*. 2021), four major haplotypes were found, which differ in the integration of transposable elements (**Fig. 5e**). Haplotype 4 only has one line, TX303. The k-mers located across the boundary of transposon insertions were detected to be associated with ULA. Tissue specific analysis showed that *MADS69* had high expression levels in the meristematic tissues, namely, ear, internode, callus, and root (**Supplementary Fig. 8a**). RNA-Seq data from four tissues of 26 NAM founders were used to examine expression of different *MADS69* haplotypes. On average, the NAM founders with haplotype 1 had lower *MADS69* expression in ears, roots, and shoots, and overall more upright leaf angle as compared to the NAM founders with haplotypes 3. Shoot expressions of *MADS69* and leaf angles of NAM founders are negatively correlated (Pearson correlation = -0.51, *p*- value < 0.01) (**Supplementary Fig. 8b**). However, *MADS69* expressions in the tassel and leaf angles are slightly positively correlated (Pearson correlation = 0.38, p-value < 0.1). The result indicates that *MADS69* expression is dynamically controlled in different tissues from distinct genetic backgrounds (**Supplementary Table 9**). Expression examination with more relevant tissue types and ectopic expression or knockouts are needed for functional confirmation and mechanistic understanding.

In the turquoise module, ULA4 (Zm00001d002425) is orthologous to the *ENLARGED FIL EXPRESSION DOMAIN 2* (*ENF2*) gene in *Arabidopsis*. Haplotype analysis of Zm00001d002425 in 26 NAM founders revealed three different haplotypes (**Fig. 6a**). The NAM founders with haplotype 1 had overall more upright ULA than those with other haplotypes (*p*-values of t-test < 0.01, **Fig. 6b**). Expression comparison showed that gene expression of haplotype 1 lines was at higher levels as compared with expression of haplotype 2 and 3 lines in all the tissues examined, namely, root, shoot, ear and tassel (**Fig. 6c**). Moreover, expression in 739 B73 RNA-seq samples showed that Zm00001d002425 had a high level of expression in meristematic tissues, such as internode, leaf primordia, and primary root (**Fig. 6d, 6e**) (Morrison *et al*. 1994; McManus and Veit 2002). In *Arabidopsis*, loss-of-function mutants of *ENF2* had abnormal gene expression along the adxial-abaxial axis in leaf development and resulted in dynamic leaf angles (Tameshige *et al*. 2013). Collectively, the results indicate that Zm00001d002425 might function in regulation of leaf angles in maize.

**Fig. 6.**
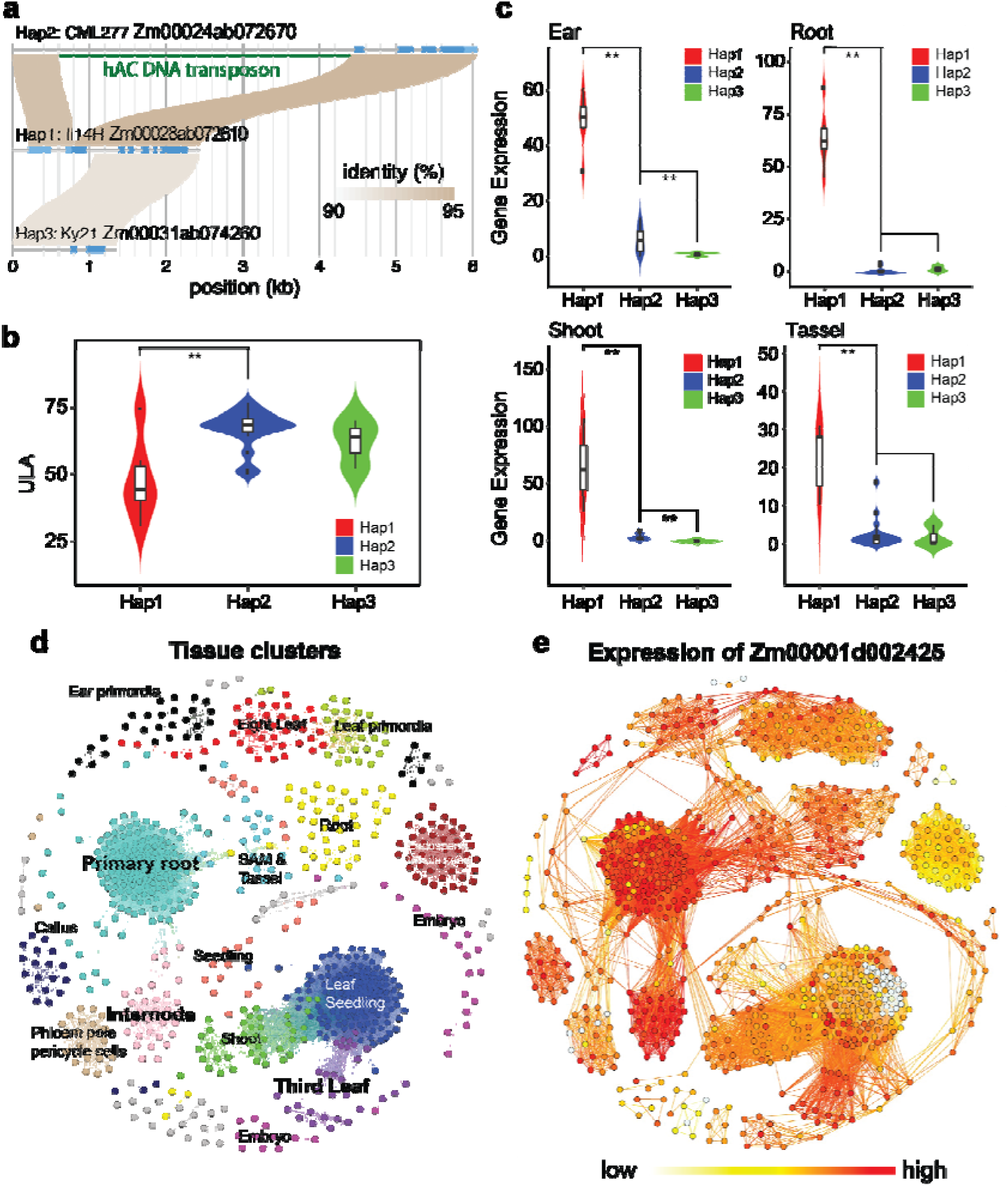
Haplotypes and expression of Zm00001d002425 (ULA4). (**a**) Alignments of three haplotypes (Hap1, 2, 3) of Zm00001d002425. Light blue and blue boxes represent untranslated regions and coding regions of the gene, respectively. The green line highlights a hAT DNA transposon in Hap2. (**b**) Violin plots for displaying distribution ULAs of NAM founders with different haplotypes. (**c**) Expressions of Zm00001d002425 with different haplotypes in multiple tissues. Double asterisks indicate that expression between two haplotypes was significantly different with a p-value less than 0.01 from a t-test. (**d**) Tissue specific expressions of Zm00001d002425 in 739 B73 RNA-Seq data. Each point represents a tissue sample. Modules are color-coded and representative tissue types are labeled. (**e**) Zm00001d002425 expression in 739 samples. The coordinate of each sample is the same as that in (d).

### K-mer GWAS of flowering time

Flowering time in maize, a trait controlled by numerous genomic loci (Buckler *et al*. 2009), was selected to demonstrate the use of k-mers for both GWAS and phenotypic prediction. K-mer GWAS on the Days To Silk (DTS) data of 259 lines from the maize282 set (**Supplementary Table 1)** (Peiffer *et al*. 2014) identified 82,678 flowering time association k-mers using the modified Bonferroni multiple test correction (p-value < 2.11e-08) (**Data S6**). Most of these k-mers (82.5%) were negatively correlated with flowering time, suggesting that the presence of most associated k-mer sequences corresponded to earlier flowering time (**Supplementary Fig. 9**).

### Genomic prediction of quantitative traits using k-mers

Several machine learning methods were developed and tested to investigate the utility of k-mer data for genomic prediction of quantitative traits. K-fold cross-validation was used to determine the accuracy of these methods in comparison to each other and standard SNP-based genomic best linear unbiased prediction (gBLUP). Randomly chosen k-mers and k-mers associated with the quantitative traits were both used for predictions. Overall, average prediction accuracies (here defined as one minus relative root mean squared error (1-rRMSE) ranged from 0.54 to 0.96 depending on the model, dataset and phenotype in question (**Fig. 7**). As expected, models designed to control for population structure performed more poorly and had greater variation in accuracy between folds than those without population structure control (**Fig. 7**). While the k-mer based methods performed slightly better than SNPs, no statistically significant differences could be detected between the best performing models with random k-mers, GWAS derived k-mers, and the SNP-based gBLUP.

**Fig. 7.**
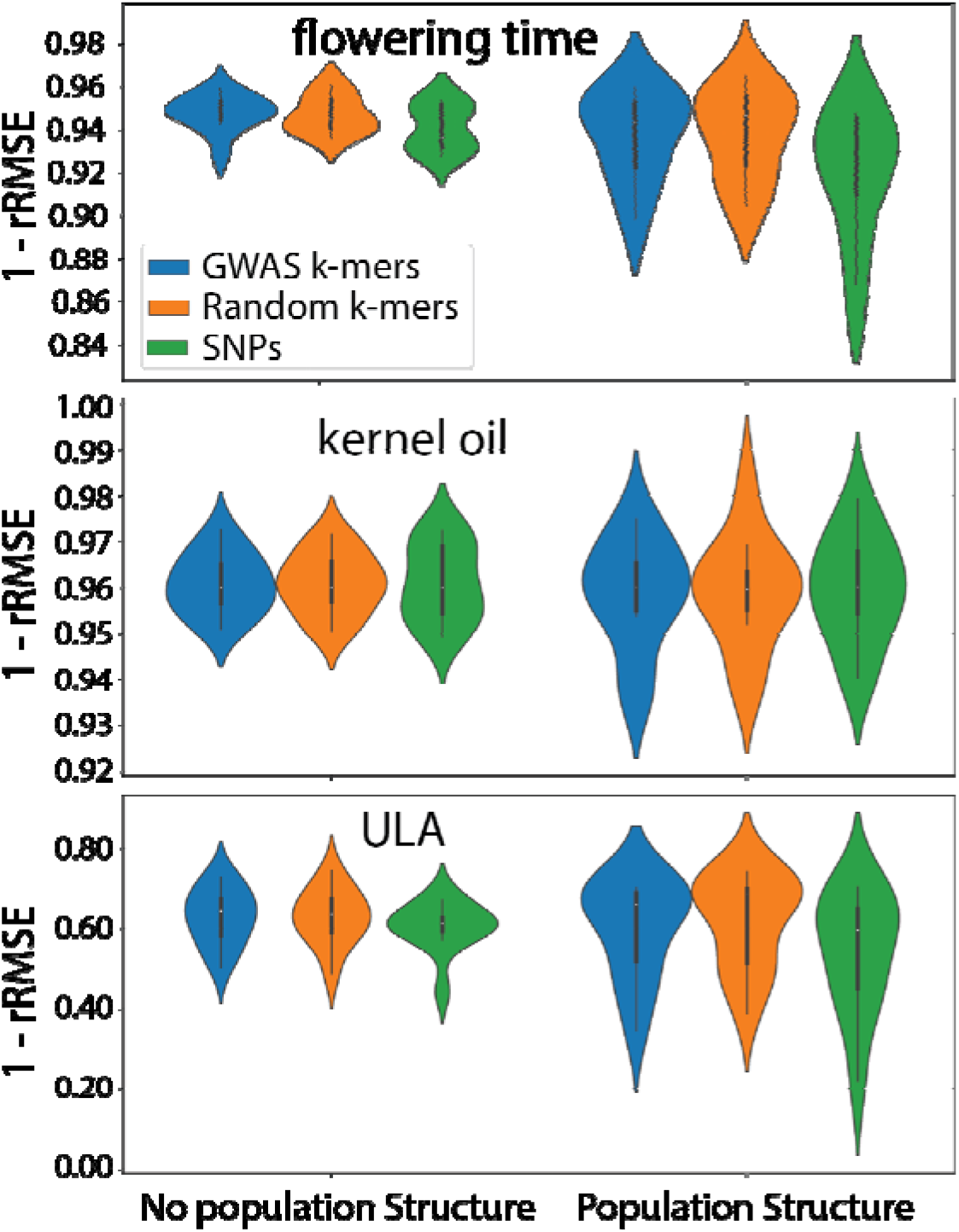
K-mer based genomic prediction. Performance of different machine learning and traditional SNP based genomic prediction methods with and without controlling for population structure for **(a)** flowering time, **(b)** kernel oil content, and **(c)** upper leaf angle. GWAS k-mers represents the best performing machine learning model when trained and tested on the top one million associated k-mers from GWAS. Random k-me is the same except that it is for the top one million randomly chosen k-mers. SNPs represent the performance of traditional SNP based genomic prediction. 1-rRMSE = one minus relative root mean squared error.

Within the machine learning models, no single method performed best across all test cases (**Supplemental Fig. 10** and **Supplemental Table 10**). When using the term frequency-inverse document frequency (TF-IDF) vectorizer, all four models performed similarly regardless of the use of copy number data or presence versus absence data. A TF-IDF vectorizer weights the values of k-mers in the model so as to lower the impact of extremely common k-mers that may overwhelm the signal and cause model overfitting. When a count vectorizer strategy was used, the neural network approach performed poorly in comparison to the other approaches, and the lasso model performed better than the other methods when copy number was included (**Supplemental Fig. 10**). These results indicate that k-mer methods can perform at least as well as standard SNP genomic prediction methods. They also suggest that the addition of k-mer copy number data to the machine learning methods here tested can result in model overfitting if not controlled for through methods such as lasso or TF-IDF.

## DISCUSSION

K-mer GWAS was employed to use WGS data from hundreds of maize lines for identifying trait-associated DNA variation. We showed the efficacy of k-mer GWAS and causal genes could be pinpointed when appropriate additional data were integrated. Both simple and complex traits were examined with k-mer GWAS and we showed that CNVs as causal variants can be directly detected. The reference-free k-mer GWAS also enables mapping of trait-associated k-mers or longer sequences assembled with associated k-mers to any available reference genomes. In addition, the simplicity of k-mers allows the integration of genomic data with expression data to reduce spurious association, and with evolutionary data to understand historic selection of genetic variants of traits.

In this study, new strategies were proposed to improve k-mer GWAS analysis. First, k-mers from sequencing reads were filtered to retain those that are present multiple times in reads of a line, measured by the KOC, from at least a portion of lines, and with variable occurrences across lines. The variability of KOCs were measured through comparison with library sizes or sequencing depths of the lines. KOCs of k-mers with equal copy numbers across genomes are expected to be highly correlated to library sizes and, therefore, were excluded in GWAS analysis. On the other hand, the total KOCs of the set of k-mers having a single copy in all genomes (i.e., conserved single-copy k-mers) served as the normalization factor to generate comparable normalized KOC data. Second, the modified Bonferroni correction considering the dependency among k-mers increases the statistical power of k-mer GWAS. The conservativeness of the method is between the Bonferroni correction and the FDR (false discovery rate) correction (Benjamini and Hochberg 1995), which enables the generation of a reasonable p-value cutoff to declare significantly associated k-mers. Third, co-occurrence networks were introduced to group k-mers based on their KOCs of the lines. Typically, physically linked k-mers can be clustered into the same group and the sequence assembly of k-mers within a cluster result in longer sequences. Sequences longer than the k-mer size can be more confidently mapped to the reference genome, which, in turn, provides clues for physical sources of all k-mers in the cluster.

Fourth, the k-mer can be a bridging element to connect various sequencing data. We integrated trait-associated k-mers with expression data to reveal the k-mers that are co-expressed with known genes in relevant pathways. The underlying assumption is that functional correlated genes tend to be co-regulated (Mao *et al*. 2009; Zeng *et al*. 2019), which implies that trait-associated k-mers co-expressed with pathway genes are more likely to be from causal variants responsible for a trait. The integration of k-mer GWAS results with expression data worked surprisingly well on the kernel color trait. Two known causal genes, *y1* and *ccd1*, were directly pinpointed with almost all the spurious associations removed. This is an ideal case but requires that some genes in the pathway are known and that most of these known genes are co-regulated. We also integrated oil and ULA k-mers with evolutionarily selective k-mers through comparing their presence in the maize population and the population of teosinte lines, the relatives to the maize ancestor. The expected historic selection during maize domestication and/or improvement confirmed the findings from k-mer GWAS. The integration with evolutionary data should reduce the level of spurious associations, thereby providing a shortened k-mer list for causal gene discovery. From the results, most alleles that benefit the kernel oil content existed in teosinte and they were largely removed to guarantee the high kernel yield in modern inbred lines. Some inbred lines, such as P39 and Oh7B, however, contained these high-oil alleles. For leaf angle, although most genomic variants conferring upright leaf angle are positively selected in maize population, some variants from either maize or teosinte populations await mining for the further optimization of leaf angle (Tian *et al*. 2019).

This study together with previous two k-mer GWAS studies has shown the power of the k-mer-based association analysis, which, in particular, is reference-free and flexible to integrate with additional data to leverage association results. Nonetheless, the approach still has room for further improvements. First, a reasonably higher depth of whole genome sequencing data from a greater number of individuals would improve statistical power, particularly for complex traits. The higher sequencing depth would reduce variation of KOCs. The average depth per line of the current study is approximately 5x. With the sequence cost continuously dropped, a higher depth of 20x or higher of each of hundreds of lines is affordable with community efforts. For simple traits, such as cob colors, 136 lines have been shown to be sufficient. But for complex traits like flowering time, additional lines would empower the detection of phenotypic associations of low-frequency alleles or with smaller effects. Second, optimized k-mer sizes and annotation of k-mers could be provided for each species. We used both 25- mers and 31-mers, in which k equals 25 and 31, respectively. Comparisons between 25-mers and 31-mers found that both gave similar results in k-mer GWAS (Voichek and Weigel 2020). The impacts of sequencing depths and quality, genome sizes, and genome complexity on the selection of k-mer sizes remains unclear. In general, smaller k-mers are more tolerant to sequencing errors and computationally more efficient, while longer k-mers are more specific and computationally more expensive (He *et al*. 2020). In addition, annotations of k-mers, such as physical sources, functional associations, and epigenomic features, could facilitate k-mer GWAS. Such annotations are possible with epigenomic data and the availability of both genome sequences and annotations of pan-genomes. Third, a dedicated statistical testing approach for k-mer GWAS is needed. The HAWK k-mer GWAS method was designed for case-control studies and population structure was accounted for (Rahman *et al*. 2018). We employed a linear regression model with controlled population structure, which proved to be effective. However, a more sophisticated approach needs to consider the handling of count data, and to control for both population stratification or structure (Q) and kinships (K), which is referred to as the QK model that is widely used in regular GWAS (Yu *et al*. 2006).

In addition to k-mer GWAS, we employed machine learning modeling for genomic prediction of quantitative traits. These models performed at least as well as standard SNP-based models and have some of the same benefits of k-mer GWAS, such as being alignment free. The use of k-mers associated with the phenotypes in k-mer GWAS did not result in greater prediction accuracy than the use of randomly sampled k-mers. This finding is not altogether surprising as the SNP-based genomic prediction literature contains mixed results on the utility of selecting SNPs for genomic prediction through GWAS (Solberg *et al*. 2008; Odilbekov *et al*. 2019). It is likely that randomly sampled k-mers (or SNPs) provide sufficient coverage of many linkage blocks for tagging genomic variation responsible for traits. Overall, the k-mer models performed very well and show promise for use in breeding and other predictive tasks.

## METHODS

### Whole Genome Sequencing (WGS) Data

WGS data consisted of 269 maize inbred lines from the Maize282 Association Panel (Bukowski *et al*. 2018). Maize282 raw sequencing data were downloaded from CyVerse (/iplant/home/shared/panzea/raw_seq_282).

### Trimming of WGS reads

Adaptor sequences and low-quality sequences on reads were trimmed with Trimmomatic (version 0.36) (Bolger *et al*. 2014) with the parameters of “ILLUMINACLIP: <adaptor>:3:20:10:1:true LEADING:3 TRAILING:3 SLIDINGWINDOW:4:13 MINLEN:50”. The adaptor database contains both TruSeq2 and TruSeq3 primer and adaptor sequences (http://www.usadellab.org/cms/?page=trimmomatic). After trimming, 8 lines were filtered out due to the low read depths and 261 lines were retained for further analysis.

### K-mer occurrence count data

Trimmed sequences were subjected to k-mer counting using the count function in Jellyfish (version 2.2.7) (Marçais and Kingsford 2011) with the k-mer size of 25 nt. K- mers with at least five read counts in at least 10 lines were retained. Occurrence counts of all these k-mers, referred to as KOC, in reads of each line were then determined.

### Normalization of KOCs across lines

KOCs were normalized to be comparable across lines. We modified the previous method that utilized a set of “conserved single-copy” k-mers for normalization (Liu *et al*. 2017). Briefly, this method first identified single-copy k-mers in both B73 and Mo17. Using WGS of HapMap2 lines, we identified a subset of single-copy k-mers showing conserved across HapMap2 lines (Hufford *et al*. 2012). The conservation was determined by finding a high correlation between KOC and library size (total reads). Here, we adjusted the procedure by taking 10% top k-mers with the highest correlation rather than top 5% using previously (Liu *et al*. 2017). And then the set was filtered by removing k-mers at 10% tails from both sides of the k-mer abundance distribution using a high-depth (∼40x) A188 sequencing dataset, and 10% tails from both sides of the total k-mer abundance distribution from all HapMap2 lines. The final single-copy conserved k-mer set (scc-kmer25.v2) contains 68,452 k-mers (**Data S5**). The total count (*C*) of all scc-kmer25.v2 k-mers of each maize282 line was determined. The normalization factor for the *i*th line was calculated by using the formula 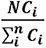, where *N* is the total number of maize282 lines. The large matrix of filtered normalized KOCs were used for k-mer GWAS.

### Removal of k-mers with low KOC variation

K-mers were further filtered if KOCs of a k-mer are highly correlated with the normalization factors. The 0.8 was used as the correlation cutoff. This step removed k-mers whose KOCs can be largely explained by sequencing depths of maize samples.

### K-mer GWAS

A linear model was used to identify trait-associated k-mers. The response variable in the model is the trait value. For the method of “counts”, the full model contains fixed effects of the normalized KOCs of a k-mer and the three top principal components derived from SNP data, representing population structure. The reduced model only contains the three principal components. Comparison of two models was performed to obtain a *p*-value for the null hypothesis that the KOCs of the k-mer are not associated with trait values. The cob color phenotypic data were collected from the Germplasm Resources Information Network (GRIN) database. The kernel color phenotypic data of 261 maize lines were measured and converted to two types: yellow and white. The oil, upper leave angle and flowering time phenotypic data were all downloaded from the Panzea dataset (https://www.panzea.org/phenotypes). Principal Components Analysis (PCA) was performed with the high-quality SNPs in Maize HapMapV3.2.1_AGPv4 dataset with Minor Allele Frequency (MAF) > 0.3 and missing rate < 0.1 (Bukowski et al., 2015). The top three components were used to represent the population structure. Three other k-mer GWAS approaches, namely, log2 counts, presence/absence, copy number, were employed. For “log2 counts”, log 2 values of normalized KOCs were used to replace normalized KOCs in the “counts” method. For “presence/absence”, all normalized KOCs were converted to either 1, if normalized KOCs are positive, or 0, if normalized KOCs equal 0. For “copy number”, the copy number of each k-mer is 0 if the normalized KOC equals 0. Otherwise, the copy number is 1 or higher depending on the rounding value of the ratio of KOC to the mean depth of the line. If the rounding value is smaller than 1, the copy number is 1. In our method, the ratio was shrunk by dividing with 2 to be conservative in calling higher copies. For example, for a k-mer, if the KOC is 9 and the mean depth is 5, the copy number is the rounding value of 9/5/2, i.e., 1.

A modified Bonferroni correction was used to account for multiple association tests. The modified method considers the dependency among k-mers. In detail, one million k-mers were randomly extracted and pairwise correlations were determined. We computed the average times of each k-mer (***m***) that was highly correlated with other k-mers (a high correlation of 0.9 was used as the threshold) based on normalized KOCs. A subset of k-mers (*N* = 10^6^) are used to reduce computational time. Based on that, the mean number of k-mer groups with a high correlation in the whole k-mer set is 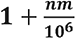, of which one is added to guarantee that at least 1 is in each group. The modified Bonferroni p-value threshold at the significant level of « is then calculated by *formula 1*. If none of the k-mer pairs are highly correlated, m is equal to 0 and the classical Bonferroni correction is performed. Any non-zero m enlarges the p-value threshold, thereby increasing the statistical power for the discovery of trait-associated k-mers. 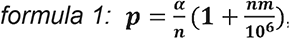, where α=0.05 and u is the total k-mers in GWAS.

### SNP GWAS

TASSEL5.0 (Bradbury *et al*. 2007) was used to perform SNP GWAS with a mixed linear model considering both population structure and kinships (Yu *et al*. 2006). SNP genotyping data were extracted from the Maize HapMapV3.2.1_AGPv4 genotypes (Bukowski *et al*. 2018). The same phenotypes and population structure data analyzed in k-mer GWAS were used in SNP GWAS. The kinship of maize lines was calculated by the GAPIT software (Lipka *et al*. 2012).

### Mapping associated k-mers directly to a reference genome

For cob colors, the associated k-mers were directly aligned to the reference genome using Bowtie (version 1.2.2) with the parameters ‘-v 2 -a -f’ (Langmead 2010). For other traits, the associated k-mers were first assembled to longer sequences with Cap3 (Huang and Madan 1999) using the parameters of “-i 25 -j 35 -s 300 -o 20 -p 94”. Assembled contigs and unassembled k-mers were then mapped to the reference using BWA (0.7.17-r1188) (Li 2010). Sequences with at least 70 bp were aligned with the “mem” module and other shorter sequences were aligned with the modules of “aln” and “samse”. All alignments were merged as final alignments.

### K-mer co-occurrence network (CON) analysis

Normalized KOCs of trait-associated k-mers in 261 lines were input to construct CONs using WGCNA (version 1.66) (Langfelder and Horvath 2008). The scale independences were determined to decide the best soft-thresholding power for analysis of network topology. Different parameters were then used to group different trait-associated k-mers into clusters by WGCNA. For the kernel color associated k-mers, the parameters were set as “power=18, TOMType = “unsigned”, minModuleSize = 150, reassignThreshold = 0, mergeCutHeight = 0.1, verbose = 3, deepSplit = 2”. For the flowering time associated k-mers, the parameters were set as “power=14, TOMType = “unsigned”, minModuleSize = 200, reassignThreshold = 0, mergeCutHeight = 0.1, verbose = 3, deepSplit = 2”. The connectivity with threshold > 0.2 was set to extract k-mers that are strongly connected. For the k-mers that failed to construct WGCNA networks, we perform the k-mean approach to cluster k-mers with the ‘kmeans’ function in R package ‘Cluster’ (Kaufman and Rousseeuw 2009). K-mers from each CON cluster were *de novo* assembled into contigs with Cap3 (Huang and Madan 1999) using the parameters of “-i 21 -j 31 -s 251 -o 16 -p 90”. Contigs were aligned to a selected reference genome through BWA. The mapping location of a cluster was determined if at least 60% mapped contigs from the cluster were uniquely mapped to a chromosomal region smaller than 10 Mb.

### K-mers from RNA-Seq reads

The same trimming and k-mer counting methods used on WGS data were performed on 739 published B73 RNA-Seq datasets (**Data S2**). K-mers presented in less than 10 samples from all datasets were removed and the remaining k-mers were normalized by RNA-Seq library sizes. KOCs of RNA-Seq reads were used as expression levels of k-mers. To identify gene expression, trimmed RNA-Seq data were aligned to the B73 reference genome (B73v4) by the STAR software (Dobin *et al*. 2013) with the gene annotation of B73v4 (Zea_mays.B73_RefGen_v4.43.gtf). Read counts per gene were determined by STAR as well. The parameters of STAR were set as “--alignIntronMax 100000 --alignMatesGapMax 100000 --outSAMattrIHstart 0 --outSAMmultNmax 1 -- outSAMstrandField intronMotif --outFilterIntronMotifs RemoveNoncanonicalUnannotated --outSAMtype BAM SortedByCoordinate -- quantMode GeneCounts --outFilterMismatchNmax 2 –outFilterMismatchNoverLmax 0.02 --outFilterMatchNmin 50 --outSJfilterReads Unique --outFilterMultimapNmax 1 -- outSAMmapqUnique 60 --outFilterMultimapScoreRange 2”. Raw read counts of each gene were normalized by total uniquely mapped reads of RNA-Seq samples.

### Construction of tissue clustering network

The normalized gene read counts of 739 B73 RNA-Seq dataset was also used to construct the tissue network with the R package WGCNA (version 1.66) ((Langfelder and Horvath 2008)). WGCNA was performed to cluster 739 B73 RNA-Seq samples with the parameters minModuleSize = 20 and soft-thresholding power = 12. If > 90% samples in one module belong to the same tissue, then the module was annotated with this tissue. The Gephi software (version 0.9.2) was used to visualize tissue networks with the module and connectivity information from the WGCNA result. With the tissue network, expression of a gene can be displayed to examine its expression pattern across tissues.

### Co-expression network analysis of k-mers and genes

Occurrence counts of trait-associated k-mers in RNA-Seq reads, representing expression of k-mers, and expression of genes in related pathways were input for constructing co-expression networks with the R package WGCNA (Langfelder and Horvath 2008). Note that trait-associated k-mers that did not overlap with RNA-Seq reads were excluded. The parameters used for co-expression network analysis were set as “TOMType = “unsigned”, minModuleSize = 100, reassignThreshold = 0, mergeCutHeight = 0.1, verbose = 3, deepSplit = 2”. The outputs of co-expression analysis were visualized with Gephi (version 0.9.2) (Bastian *et al*. 2009).

### QTL analysis in the NAM population

Genotype and phenotype data of NAM recombinant inbred lines (RILs) were collected from previous studies (Glaubitz *et al*. 2014). The R package ‘qtl’ was used to calculate the LOD for each marker in each NAM RIL subpopulation. The LOD threshold was determined by performing 1,000 times permutation. For each peak, the marker with the highest LOD value was used to represent the QTL peak.

### Expression of candidate genes in NAM founders

RNA-Seq data of four tissues from NAM founders were downloaded from Sequence Read Archive (SRX129687). Data processing, including read trimming, alignment, and read counts, was similar to the process for analyzing 739 B73 RNA-Seq.

### Identification of evolutionarily selected k-mers

WGS data of 20 teosintes were collected from a previous study (Xu *et al*. 2020). The same methods were performed to generate the k-mers and KOCs from them. To identify evolutionarily selected k-mers, all trait associated k-mers were examined through aχ^2^ statistical test between 20 teosintes and 257 inbred lines. The hypothesis of the test is that the proportion of lines carrying a given k-mer in teosinte lines is not different from that in maize282 lines. Multiple tests were accounted for to control the false discovery rate at 5% (Benjamini and Hochberg 1995). k-mers with the significance were considered as evolutionarily selected k-mers. Frequencies of each of these k-mers in teosinte or maize lines were calculated by counting the proportion of lines with the k-mer (KOC≥1).

### Prediction of quantitative traits

Multiple machine learning strategies were implemented and tested for predicting the flowering time, kernel oil, and upper leaf angle phenotype from k-mer data. These models relied strongly on variations of NLP methods, particularly “bag-of-words” methods (Mejía-Guerra and Buckler 2019). KOCs were vectorized into matrices using either the CountVectorizer or TfidfVectorizer methods in the scikit-learn package (Van Rossum and Drake 2009; Pedregosa *et al*. 2011). The TfidfVectorization method was also supplemented with code and methods adapted from a previous publication (Mejía-Guerra and Buckler 2019). Various common regression frameworks were used to train the models. These methods were also implemented using the scikit-learn package and included: Linear regression, Ridge regression, and Lasso. Additionally, a shallow neural network model was designed and implemented for phenotype prediction using Keras (Chollet 2015). In addition to the machine learning methods, a standard SNP-based prediction was implemented for comparison. Briefly, genotypic data of the maize diversity panel was obtained from the Panzea database (Bukowski *et al*. 2018). By pruning SNPs in LD (*r*^2^ ≥ 0.1), a set of ∼800,000 SNPs (MAF >= 0.01) was retained using the PLINK1.9 software (Chang *et al*. 2015). The genome-enabled prediction was conducted using the “BayesC” method of GenSel v4.1 (Habier *et al*. 2011) following a two-step model training procedure as described previously (Yang *et al*. 2018).

Each of the prediction methods described above was trained/calibrated and tested using two different k-mer data sets: one million k-mers with the lowest p-values from k-mer GWAS of a corresponding trait, and a set of one million random k-mers from the GWAS analysis. A k-folds strategy was used to train and test all of the k-mer and SNP-based prediction models. K-Means clustering was used to divide the inbred lines into 11 clusters based on population structure. The number of clusters was chosen visually from a graph of possible k values and their corresponding Sum of Squared Distances (SSD). These clusters were used to test the models’ performances for predicting across distinct genetic groups. A matching set of 11 random clusters was used to test prediction performance without controlling for population structure. Pearson correlation coefficients (r), Root Mean Squared Error (RMSE), relative RMSE (rRMSE), and 1-rRMSE were calculated on the test sets from each fold individually and averaged across folds to provide the final values for model comparisons.

## ACKNOWLEDGMENTS

Funding was provided by the USDA’s National Institute of Food and Agriculture (award no. 2018-67013-28511 and 2021-67013-35724) and by the US National Science Foundation (awards No. 1741090 and 2011500). We thank Dr. Josh Strable for the valuable discussion. This is a contribution 22-114-J from the Kansas Agricultural Experiment Station, Manhattan, Kansas.

## DATA AVAILABILITY

All related scripts have been deposited at GitHub (github.com/PlantG3/ZmKmerGWAS).

## AUTHOR CONTRIBUTION

CH, JW, and SL conceptualized the approaches of k-mer GWAS and k-mer prediction. CH, YH, ZZ, and SL developed k-mer GWAS approaches and conducted k-mer GWAS analysis; JY performed phenotypic prediction with SNPs; and JW performed phenotypic prediction with k-mers. CH, JW, and SL drafted the manuscript. All authors reviewed and revised the manuscript.

## REFERENCES

1. Abou-Deif M. H., B. B. Mekki, E. A. H. Mostafa, R. M. Esmail, and S. A. M. Khattab, 2012 The genetic relationship between proteins, oil and grain yield in some maize hybrids. World J. Agric. Sci. 8: 43–50.

2. Anvar S. Y., L. Khachatryan, M. Vermaat, M. van Galen, I. Pulyakhina, et al., 2014 Determining the quality and complexity of next-generation sequencing data without a reference genome. Genome Biol. 15: 555.

3. Ariyanayagam, R.P., Moore, C.L. and Carangal, V.R., 1974 Selection for Leaf Angle in Maize and Its Effect on Grain Yield and Other Characters. Crop Science, 14: 551–556.

4. Azmach G., A. Menkir, C. Spillane, and M. Gedil, 2018 Genetic loci controlling carotenoid biosynthesis in diverse tropical maize lines. G3 8: 1049–1065.

5. Balding D. J., 2006 A tutorial on statistical methods for population association studies. Nat. Rev. Genet. 7: 781–791.

6. Bastian M., S. Heymann, M. Jacomy, and Others, 2009 Gephi: an open source software for exploring and manipulating networks. Icwsm 8: 361–362.

7. Benjamini Y., and Y. Hochberg, 1995 Controlling the false discovery rate: a practical and powerful approach to multiple testing. J. R. Stat. Soc. Series B Stat. Methodol. 57: 289–300.

8. Bolger A. M., M. Lohse, and B. Usadel, 2014 Trimmomatic: a flexible trimmer for Illumina sequence data. Bioinformatics 30: 2114–2120.

9. Bradbury P. J., Z. Zhang, D. E. Kroon, T. M. Casstevens, Y. Ramdoss, et al., 2007 TASSEL: software for association mapping of complex traits in diverse samples. Bioinformatics 23: 2633–2635.

10. Buckler E. S., J. B. Holland, P. J. Bradbury, C. B. Acharya, P. J. Brown, et al., 2009 The genetic architecture of maize flowering time. Science 325: 714–718.

11. Buckner B., T. L. Kelson, and D. S. Robertson, 1990 Cloning of the y1 locus of maize, a gene involved in the biosynthesis of carotenoids. Plant Cell 2: 867–876.

12. Bukowski R., X. Guo, Y. Lu, C. Zou, B. He, et al., 2018 Construction of the third-generation Zea mays haplotype map. Gigascience 7: 1–12.

13. Chai Y., X. Hao, X. Yang, W. B. Allen, J. Li, et al., 2012 Validation of DGAT1-2 polymorphisms associated with oil content and development of functional markers for molecular breeding of high-oil maize. Mol. Breed. 29: 939–949.

14. Chang C. C., C. C. Chow, L. C. Tellier, S. Vattikuti, S. M. Purcell, et al., 2015 Second-generation PLINK: rising to the challenge of larger and richer datasets. GigaScience 4: 7.

15. Chollet F., 2015 Keras, https://github.com/keras-team/keras

16. Dobin A., C. A. Davis, F. Schlesinger, J. Drenkow, C. Zaleski, et al., 2013 STAR: ultrafast universal RNA-seq aligner. Bioinformatics 29: 15–21.

17. Duncan W. G., 1971 Leaf angles, leaf area, and canopy photosynthesis. Crop Sci. 11: 482–485.

18. Flint-Garcia S. A., A.-C. Thuillet, J. Yu, G. Pressoir, S. M. Romero, et al., 2005 Maize association population: a high-resolution platform for quantitative trait locus dissection. Plant J 44: 1054–1064.

19. Glaubitz J. C., T. M. Casstevens, F. Lu, J. Harriman, R. J. Elshire, et al., 2014 TASSEL-GBS: a high capacity genotyping by sequencing analysis pipeline. PLoS One 9: e90346.

20. Grotewold E., B. J. Drummond, B. Bowen, and T. Peterson, 1994 The myb-homologous P gene controls phlobaphene pigmentation in maize floral organs by directly activating a flavonoid biosynthetic gene subset. Cell 76: 543–553.

21. Habier D., R. L. Fernando, K. Kizilkaya, and D. J. Garrick, 2011 Extension of the bayesian alphabet for genomic selection. BMC Bioinformatics 12: 186.

22. He C., G. Lin, H. Wei, H. Tang, F. F. White, et al., 2020 Factorial estimating assembly base errors using k-mer abundance difference (KAD) between short reads and genome assembled sequences. NAR Genomics and Bioinformatics 2:lqaa075.

23. Huang X., and A. Madan, 1999 CAP3: A DNA sequence assembly program. Genome Res. 9: 868–877.

24. Hufford M. B., X. Xu, J. van Heerwaarden, T. Pyhäjärvi, J.-M. Chia, et al., 2012 Comparative population genomics of maize domestication and improvement. Nat. Genet. 44: 808–811.

25. Hufford M. B., A. S. Seetharam, M. R. Woodhouse, K. M. Chougule, S. Ou, et al., 2021 De novo assembly, annotation, and comparative analysis of 26 diverse maize genomes. Science 373: 655–662.

26. Jiao Y., P. Peluso, J. Shi, T. Liang, M. C. Stitzer, et al., 2017 Improved maize reference genome with single-molecule technologies. Nature 546: 524–527.

27. Kaufman, L. and Rousseeuw, P.J., 2009. Finding groups in data: an introduction to cluster analysis (Vol 344). John Wiley & Sons.

28. Langfelder P., and S. Horvath, 2008 WGCNA: an R package for weighted correlation network analysis. BMC Bioinformatics 9: 559.

29. Langmead B., 2010 Aligning short sequencing reads with Bowtie. Curr. Protoc. Bioinformatics 32: 11.7.1–11.7.14.

30. Li H., and R. Durbin, 2010 Fast and accurate long-read alignment with Burrows– Wheeler transform. Bioinformatics 26: 589–595.

31. Li H., Z. Peng, X. Yang, W. Wang, J. Fu, et al., 2013 Genome-wide association study dissects the genetic architecture of oil biosynthesis in maize kernels. Nat. Genet. 45: 43–50.

32. Li X., P. Wu, Y. Lu, S. Guo, Z. Zhong, et al., 2020 Synergistic interaction of phytohormones in determining leaf angle in crops. Int. J. Mol. Sci. 21: 5052.

33. Liang Y., Q. Liu, X. Wang, C. Huang, G. Xu, et al., 2019 ZmMADS69 functions as a flowering activator through the ZmRap2.7-ZCN8 regulatory module and contributes to maize flowering time adaptation. New Phytol. 221: 2335–2347.

34. Lin G., C. He, J. Zheng, D.-H. Koo, H. Le, et al., 2021 Chromosome-level genome assembly of a regenerable maize inbred line A188. Genome Biol. 22: 175.

35. Lipka A. E., F. Tian, Q. Wang, J. Peiffer, M. Li, et al., 2012 GAPIT: genome association and prediction integrated tool. Bioinformatics 28: 2397–2399.

36. Liu B., Y. Shi, J. Yuan, X. Hu, H. Zhang, et al., 2013 Estimation of genomic characteristics by analyzing k-mer frequency in de novo genome projects. arXiv. 1308.2012

37. Liu S., J. Zheng, P. Migeon, J. Ren, Y. Hu, et al., 2017 Unbiased k-mer analysis reveals changes in copy number of highly repetitive sequences during maize domestication and improvement. Sci. Rep. 7: 42444.

38. Liu H.-J., and J. Yan, 2019 Crop genome-wide association study: a harvest of biological relevance. Plant J. 97: 8–18.

39. Liu K., J. Cao, K. Yu, X. Liu, Y. Gao, et al., 2019 Wheat TaSPL8 modulates leaf angle through auxin and brassinosteroid signaling. Plant Physiol. 181: 179–194.

40. Lu F., M. C. Romay, J. C. Glaubitz, P. J. Bradbury, R. J. Elshire, et al., 2015 High-resolution genetic mapping of maize pan-genome sequence anchors. Nat. Commun. 6: 6914.

41. Luo X., J. Zheng, R. Huang, Y. Huang, H. Wang, et al., 2016 Phytohormones signaling and crosstalk regulating leaf angle in rice. Plant Cell Rep. 35: 2423–2433.

42. Mao L., J. L. Van Hemert, S. Dash, and J. A. Dickerson, 2009 Arabidopsis gene co-expression network and its functional modules. BMC Bioinformatics 10: 346.

43. Marçais G., and C. Kingsford, 2011 A fast, lock-free approach for efficient parallel counting of occurrences of k-mers. Bioinformatics 27: 764–770.

44. McManus M. T., and B. E. Veit, 2002 Meristematic tissues in plant growth and development. Sheffield Academic Press: 172–212

45. Mejía-Guerra M. K., and E. S. Buckler, 2019 A k-mer grammar analysis to uncover maize regulatory architecture. BMC Plant Biol. 19: 103.

46. Morrison T. A., J. R. Kessler, and D. R. Buxton, 1994 Maize internode elongation patterns. Crop Sci. 34: 1055–1060.

47. Muehlbauer G. J., J. E. Fowler, and M. Freeling, 1997 Sectors expressing the homeobox gene liguleless3 implicate a time-dependent mechanism for cell fate acquisition along the proximal-distal axis of the maize leaf. Development 124: 5097–5106.

48. Odilbekov F., R. Armoniené, A. Koc, J. Svensson, and A. Chawade, 2019 GWAS- assisted genomic prediction to predict resistance to Septoria tritici blotch in Nordic winter wheat at seedling stage. Front. Genet. 10: 1224.

49. Owens B. F., A. E. Lipka, M. Magallanes-Lundback, T. Tiede, C. H. Diepenbrock, et al., 2014 A foundation for provitamin A biofortification of maize: genome-wide association and genomic prediction models of carotenoid levels. Genetics 198: 1699–1716.

50. Pedregosa F., G. Varoquaux, A. Gramfort, V. Michel, B. Thirion, et al., 2011 Scikit-learn: machine learning in Python. J. Mach. Learn. Res. 12: 2825–2830.

51. Peiffer J. A., M. C. Romay, M. A. Gore, S. A. Flint-Garcia, Z. Zhang, et al., 2014 The genetic architecture of maize height. Genetics 196: 1337–1356.

52. Pendleton J. W., G. E. Smith, S. R. Winter, and T. J. Johnston, 1968 Field investigations of the relationships of leaf angle in corn ( Zea mays L.) to grain yield and apparent photosynthesis. Agron. J. 60: 422–424.

53. Pepper, G.E., Pearce, R.B. and Mock, J.J., 1977. Leaf orientation and yield of maize. Crop Science, 17: 883–886.

54. Rahman A., I. Hallgrímsdóttir, M. Eisen, and L. Pachter, 2018 Association mapping from sequencing reads using k-mers. Elife 7: e32920.

55. Schnable P. S., D. Ware, R. S. Fulton, J. C. Stein, F. Wei, et al., 2009 The B73 maize genome: complexity, diversity, and dynamics. Science 326: 1112–1115.

56. Simpson J. T., 2014 Exploring genome characteristics and sequence quality without a reference. Bioinformatics 30: 1228–1235.

57. Solberg T. R., A. K. Sonesson, J. A. Woolliams, and T. H. E. Meuwissen, 2008 Genomic selection using different marker types and densities. J. Anim. Sci. 86: 2447–2454.

58. Sun H., J. Ding, M. Piednoël, and K. Schneeberger, 2018 findGSE: estimating genome size variation within human and Arabidopsis using k-mer frequencies. Bioinformatics 34: 550–557.

59. Suwarno W. B., K. V. Pixley, N. Palacios-Rojas, S. M. Kaeppler, and R. Babu, 2015 Genome-wide association analysis reveals new targets for carotenoid biofortification in maize. Theor. Appl. Genet. 128: 851–864.

60. Tam V., N. Patel, M. Turcotte, Y. Bossé, G. Paré, et al., 2019 Benefits and limitations of genome-wide association studies. Nat. Rev. Genet. 20: 467–484.

61. Tameshige T., H. Fujita, K. Watanabe, K. Toyokura, M. Kondo, et al., 2013 Pattern dynamics in adaxial-abaxial specific gene expression are modulated by a plastid retrograde signal during Arabidopsis thaliana leaf development. PLoS Genet. 9: e1003655.

62. Tan B.-C., J.-C. Guan, S. Ding, S. Wu, J. W. Saunders, et al., 2017 Structure and origin of the white cap locus and its role in evolution of grain color in maize. Genetics 206: 135–150.

63. Tian J., C. Wang, J. Xia, L. Wu, G. Xu, et al., 2019 Teosinte ligule allele narrows plant architecture and enhances high-density maize yields. Science 365: 658–664.

64. Tian D., P. Wang, B. Tang, X. Teng, C. Li, et al., 2020 GWAS Atlas: a curated resource of genome-wide variant-trait associations in plants and animals. Nucleic Acids Res. 48: D927–D932.

65. Van Rossum G., and F. L. Drake, 2009 Python 3 Reference Manual. CreateSpace Independent Publishing Platform.

66. Visscher P. M., N. R. Wray, Q. Zhang, P. Sklar, M. I. McCarthy, et al., 2017 10 years of GWAS discovery: biology, function, and translation. Am. J. Hum. Genet. 101: 5–22.

67. Voichek Y., and D. Weigel, 2020 Identifying genetic variants underlying phenotypic variation in plants without complete genomes. Nature Genetics 52: 534–540.

68. Vurture G. W., F. J. Sedlazeck, M. Nattestad, C. J. Underwood, H. Fang, et al., 2017 GenomeScope: fast reference-free genome profiling from short reads. Bioinformatics 33: 2202–2204.

69. Washburn J. D., M. K. Mejia-Guerra, G. Ramstein, K. A. Kremling, R. Valluru, et al., 2019 Evolutionarily informed deep learning methods for predicting relative transcript abundance from DNA sequence. Proc. Natl. Acad. Sci. U. S. A. 116: 5542–5549.

70. Washburn J. D., M. B. Burch, and J. A. Valdes Franco, 2020 Predictive breeding for maize: Making use of molecular phenotypes, machine learning, and physiological crop models. Crop Science 60: 622–638.

71. Washburn J. D., E. Cimen, G. Ramstein, T. Reeves, P. O’Briant, et al., 2021 Predicting phenotypes from genetic, environment, management, and historical data using CNNs. Theor. Appl. Genet. 134: 3997–4011.

72. Xu G., J. Lyu, Q. Li, H. Liu, D. Wang, et al., 2020 Evolutionary and functional genomics of DNA methylation in maize domestication and improvement. Nat. Commun. 11: 5539.

73. Yang J., C.-T. E. Yeh, R. K. Ramamurthy, X. Qi, R. L. Fernando, et al., 2018 Empirical Comparisons of Different Statistical Models To Identify and Validate Kernel Row Number-Associated Variants from Structured Multi-parent Mapping Populations of Maize. G3 8: 3567–3575.

74. Yu J., G. Pressoir, W. H. Briggs, I. Vroh Bi, M. Yamasaki, et al., 2006 A unified mixed-model method for association mapping that accounts for multiple levels of relatedness. Nat. Genet. 38: 203–208.

75. Zanten M. van, T. L. Pons, J. A. M. Janssen, L. A. C. J. Voesenek, and A. J. M. Peeters, 2010 On the relevance and control of leaf angle. CRC Crit. Rev. Plant Sci. 29: 300–316.

76. J Zheng, C He, Y Qin, G Lin, WD Park et al., 2019 Co expression analysis aids in the identification of genes in the cuticular wax pathway in maize. Plant J. 97: 530–542.

77. Zhang F., and T. Peterson, 2005 Comparisons of maize pericarp color1 alleles reveal paralogous gene recombination and an organ-specific enhancer region. Plant Cell 17: 903–914.

78. Zheng P., W. B. Allen, K. Roesler, M. E. Williams, S. Zhang, et al., 2008 A phenylalanine in DGAT is a key determinant of oil content and composition in maize. Nat. Genet. 40: 367–372.

