## Supplementary Figures for "Trait Association and Prediction Through Integrative K-mer Analysis"

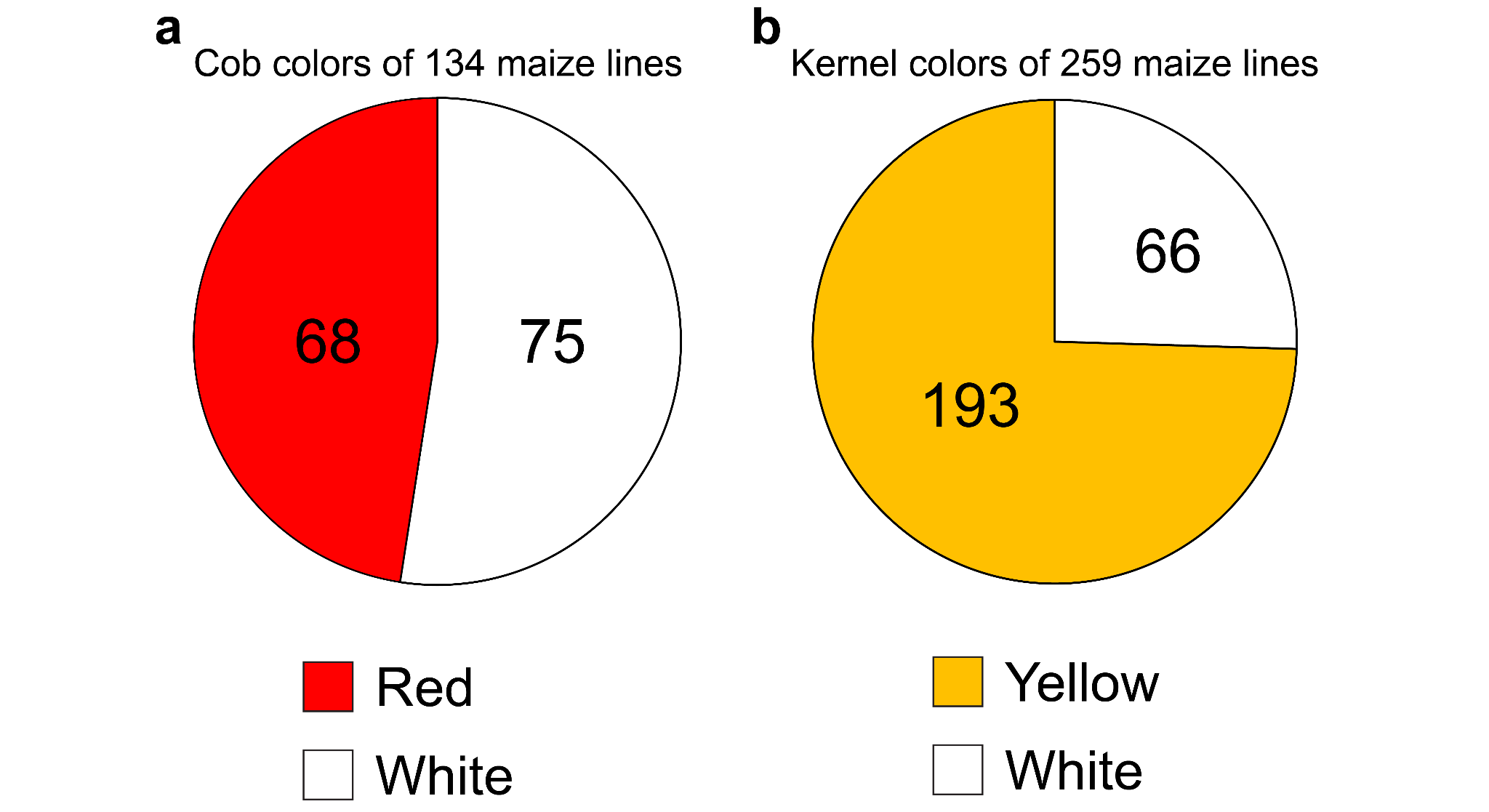

Supplementary Fig. 1. Cob color and kernel color traits of maize282. (a) Cob colors of 143 lines in the 282 maize panel. (b) Kernel colors of 259 lines in the 282 maize panel.

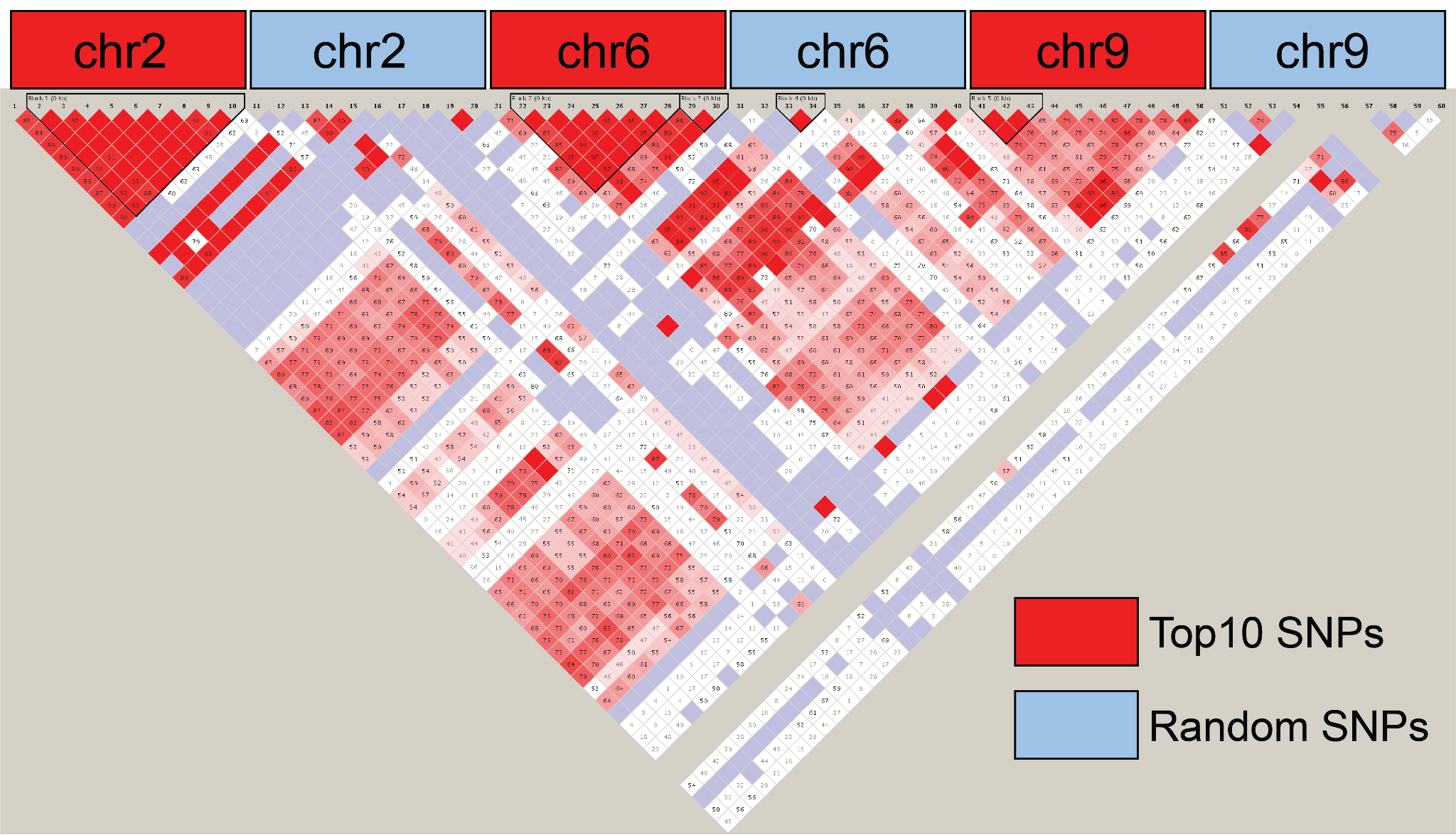

Supplementary Fig. 2. LD of SNPs from GWAS peaks of kernel color. LD analysis was performed by Haploview v4.2. The top10 SNPs from each peak on chromosomes 2, 6, and 9 and randomly selected SNPs from these three chromosomes. Colors from purple to red represent LDs from low to high.

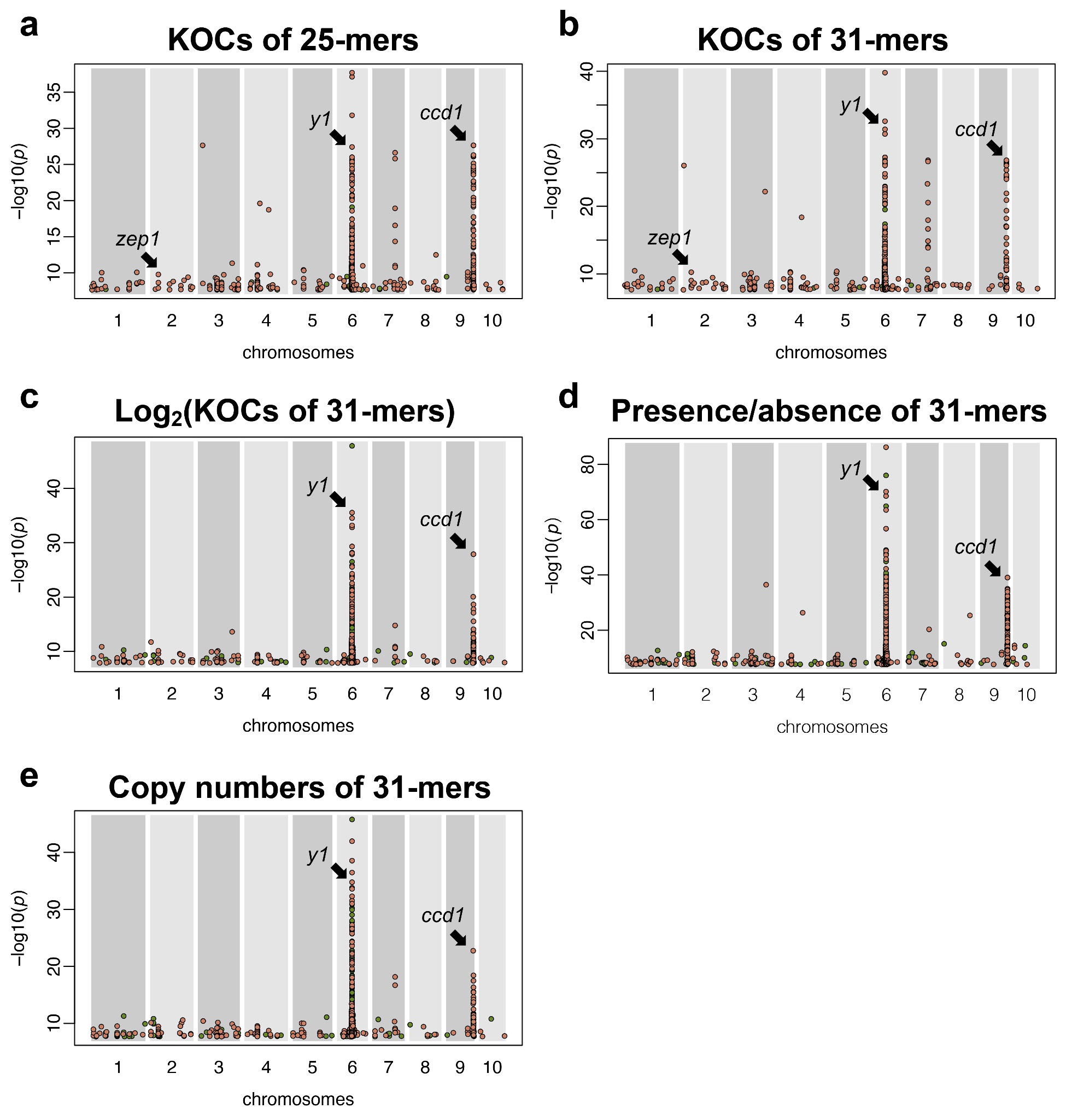

Supplementary Fig. 3. Comparison of kernel color GWAS with various methods. K-mer GWAS based on KOCs of 25-mers (a), KOCs of 31-mers (b), log2 of 31-mers KOCs (c), presence/absence of 31-mers (d), and copy numbers of 31-mers (e). The presence/absence approach had a higher power but the number of k-mers on the *ccd1* locus is much lower than other methods.

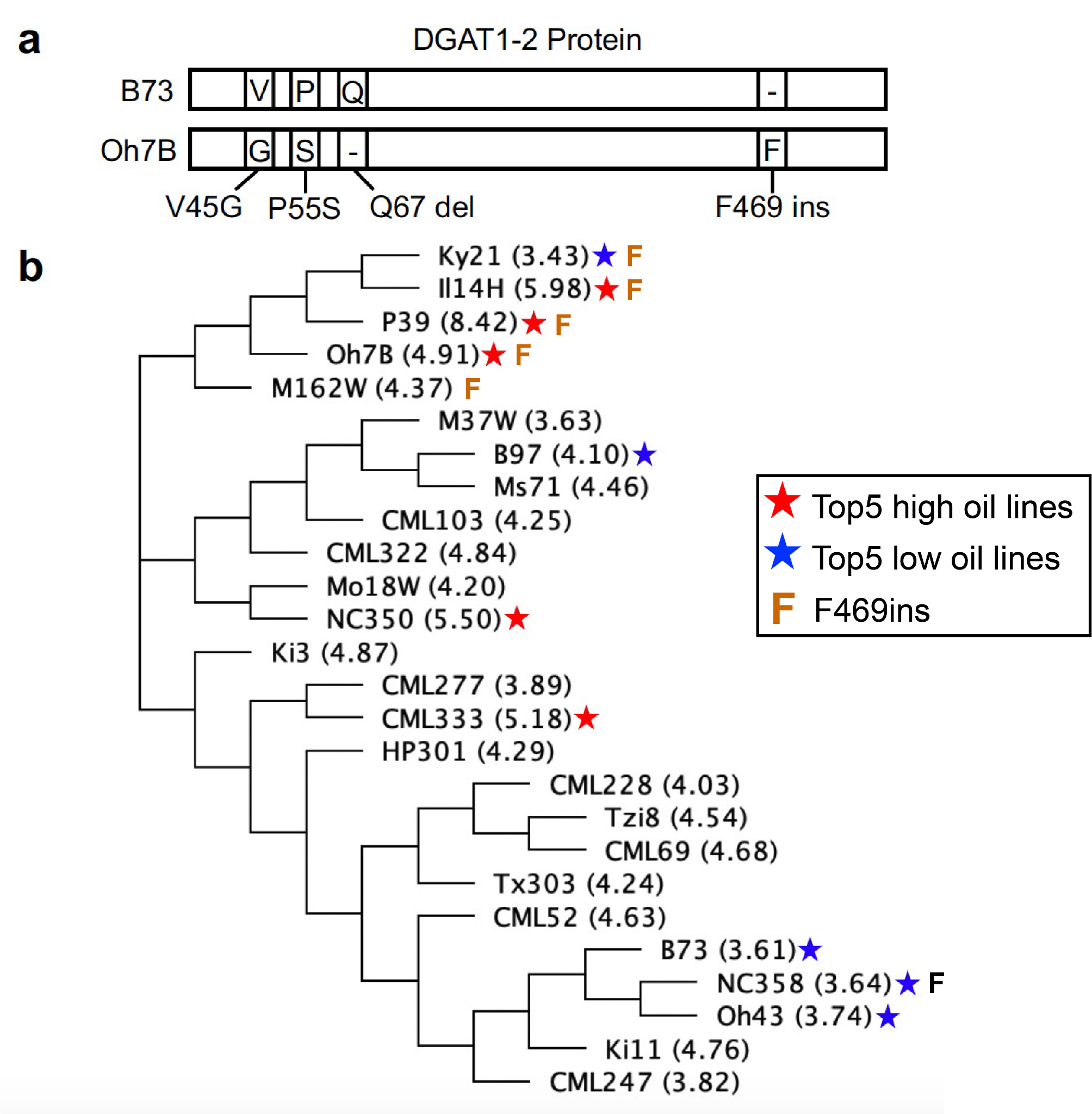

Supplementary Fig. 4. DGAT1-2 in NAMs. (a) Polymorphisms of DGAT1-2 protein sequences between B73 and Oh7B. (b) A phylogenetic tree of *DGAT1-2* gene sequences from NAM founders. Red and blue asterisks indicate lines with the highest and lowest oil contents, respectively. The “F” letter labels lines carrying F469ins.

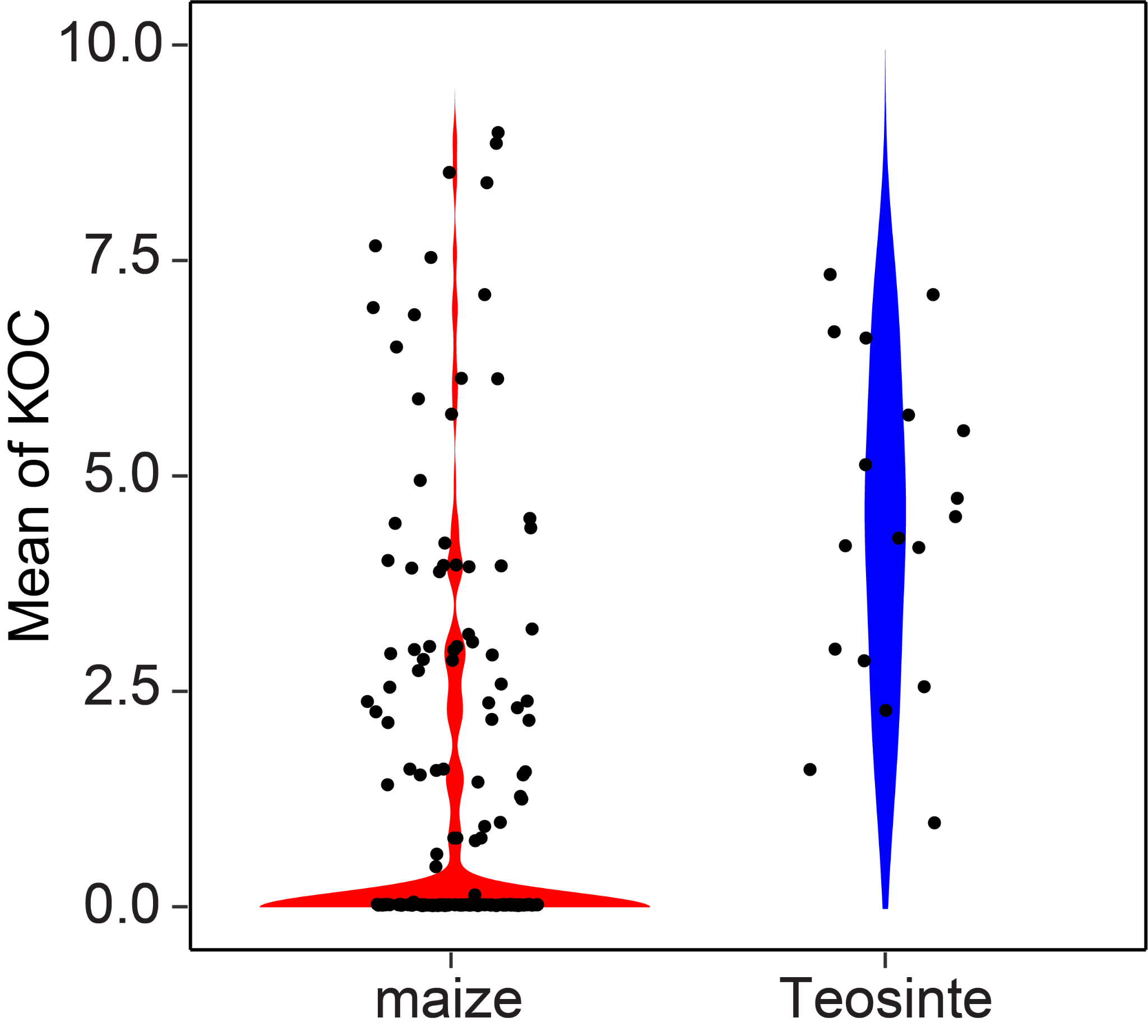

Supplementary Fig. 5. KOCs of k-mers from the F469ins region in maize and teosinte lines. The y-axis represents the mean values of normalized KOCs of all k-mers from the F469ins region per line. Each dot stands for a maize or teosinte line.

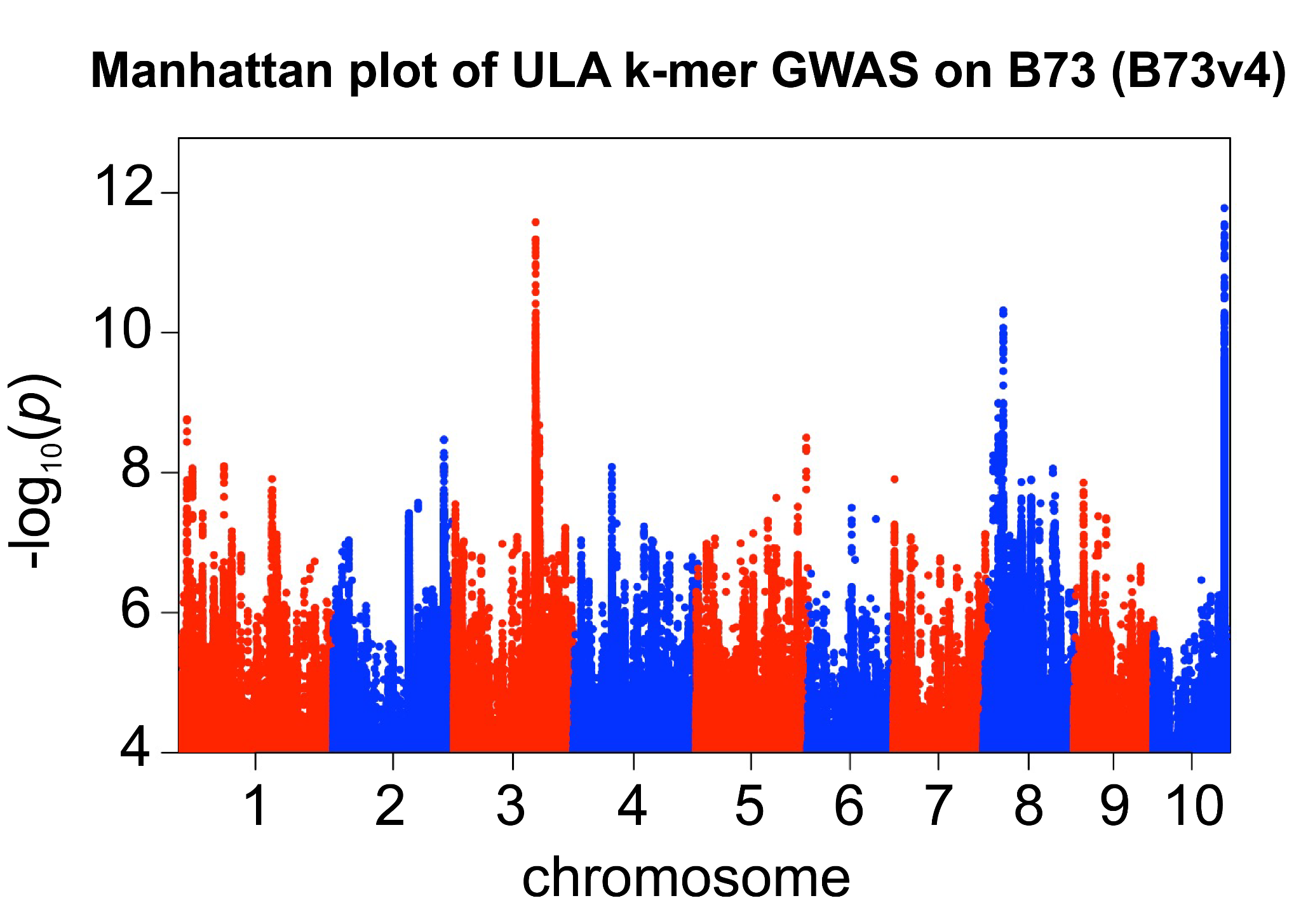

Supplementary Fig. 6. Manhattan plot of ULA k-mer GWAS on B73 (B73v4) without controlling the chromosome 10 peak. A k-mer was plotted only if it was uniquely and perfectly mapped to B73v4 and had the *p*-values less than 1e-4 from the ULA k-mer GWAS.

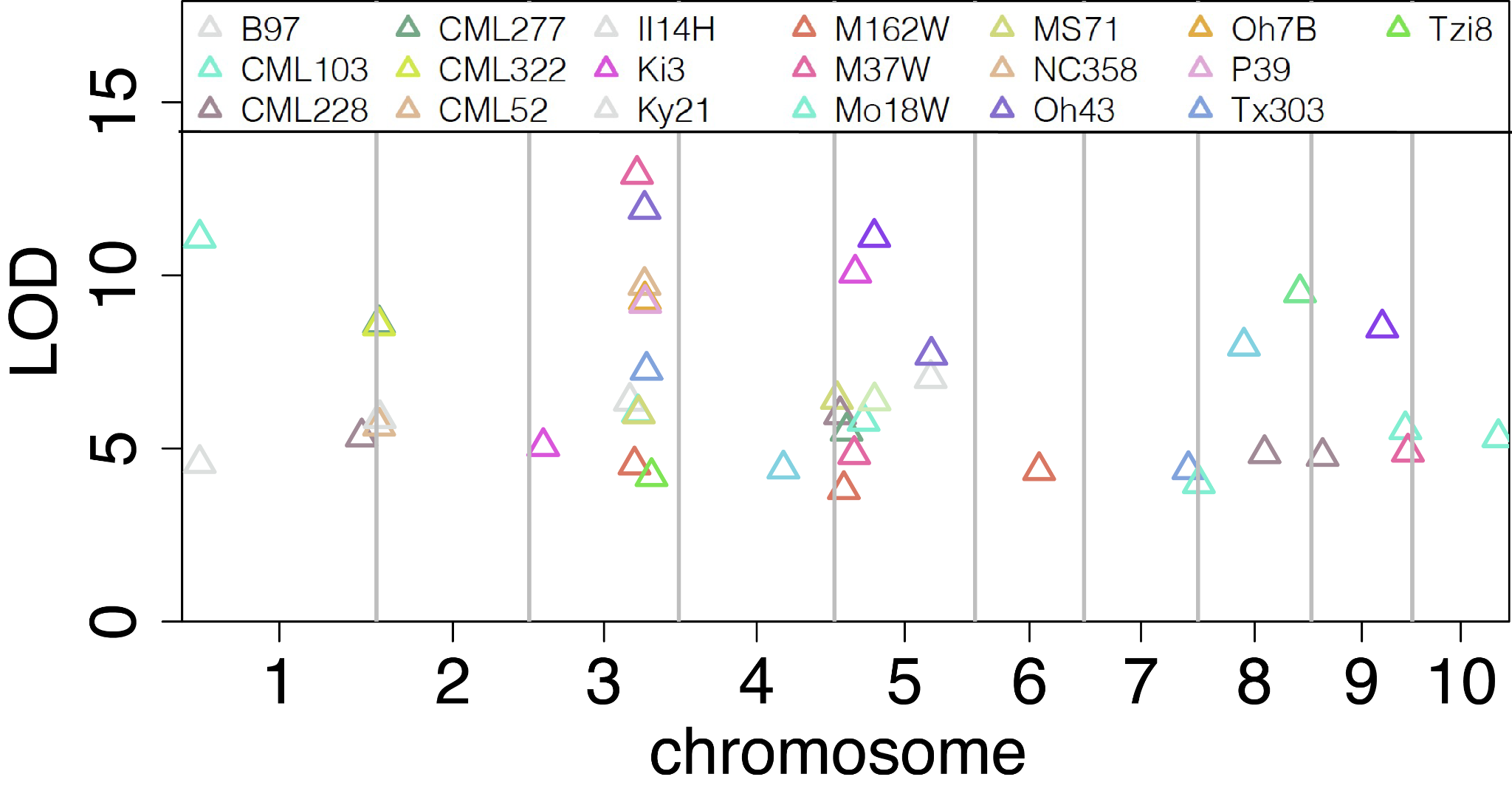

Supplementary Fig. 7. QTLs of ULA in NAM subpopulations. QTLs in different NAM subpopulations are color-coded. A marker with the highest LOD of each QTL was plotted.

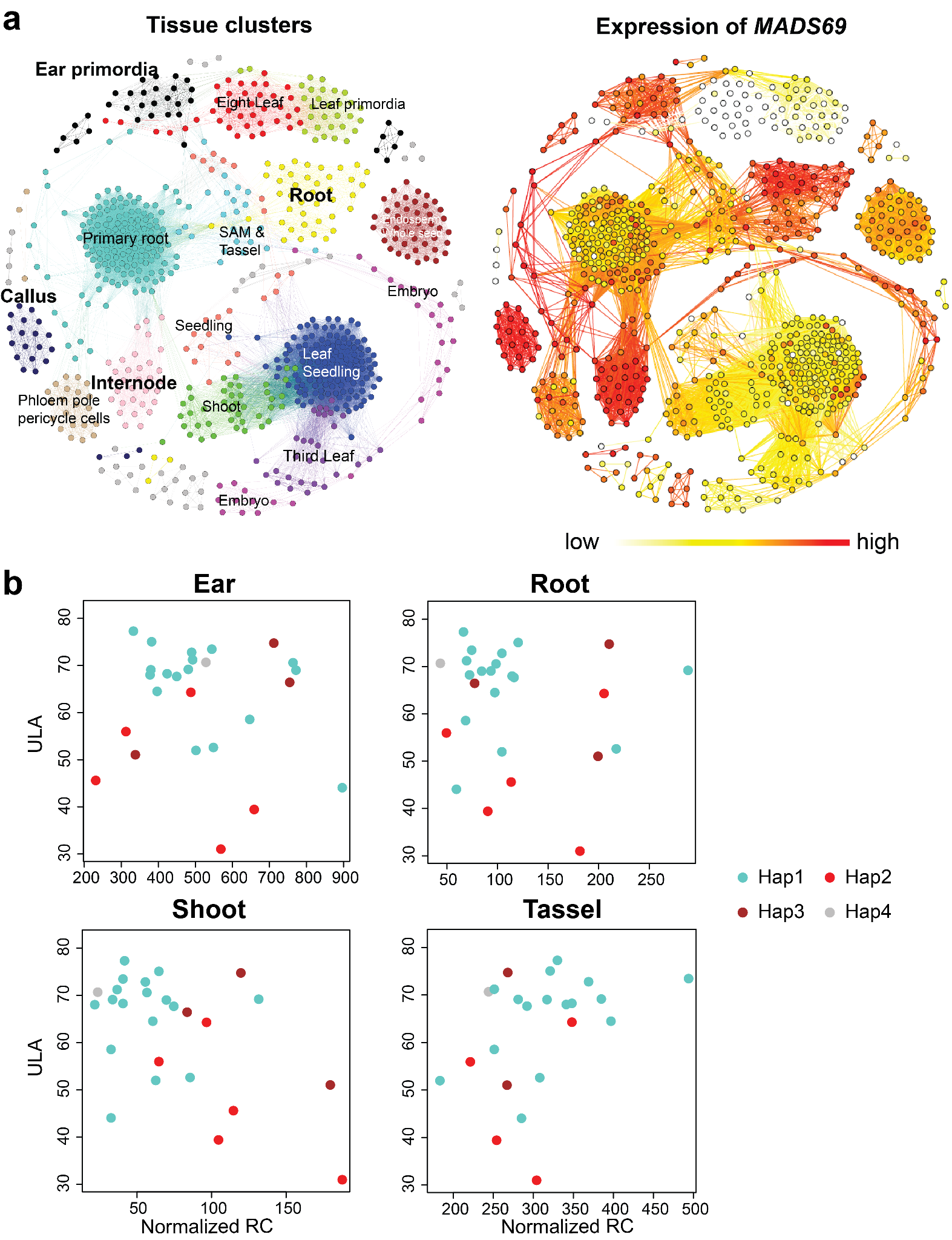

Supplementary Fig. 8. Expression analysis of Zm00001d042315 (*MADS69)*. (a) Tissue specific expressions of *MADS69* in 739 B73 RNA-Seq data. Each point represents a tissue sample. Modules are color-coded and representative tissue types are labeled. (b) *MADS69* expression in 739 tissues. The coordinate of each tissue is the same as that in (a). (c) The relationship of *MADS69* expressions and ULA traits in different tissues. The x-axis represents normalized read counts (RC) of *MADS69* and the y-axis represents the ULA phenotype. Dots with different colors represent NAM founders with different haplotypes of *MADS69*.

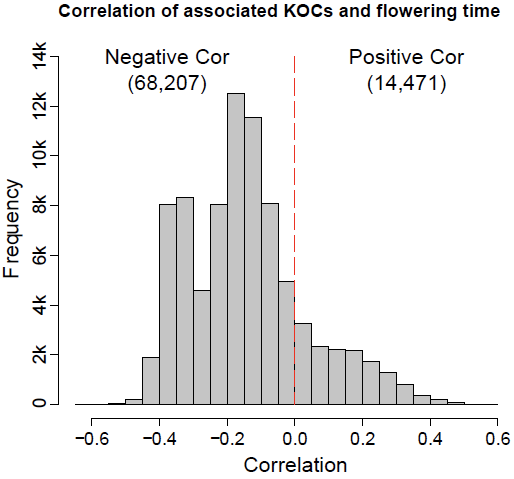
`

Supplementary Fig. 9. Correlations of KOCs of DTS associated k-mers and flowering time phenotype data. The x-axis represents the Pearson correlations of KOCs and flowering times and the y-axis represents the number of DTS k-mers.

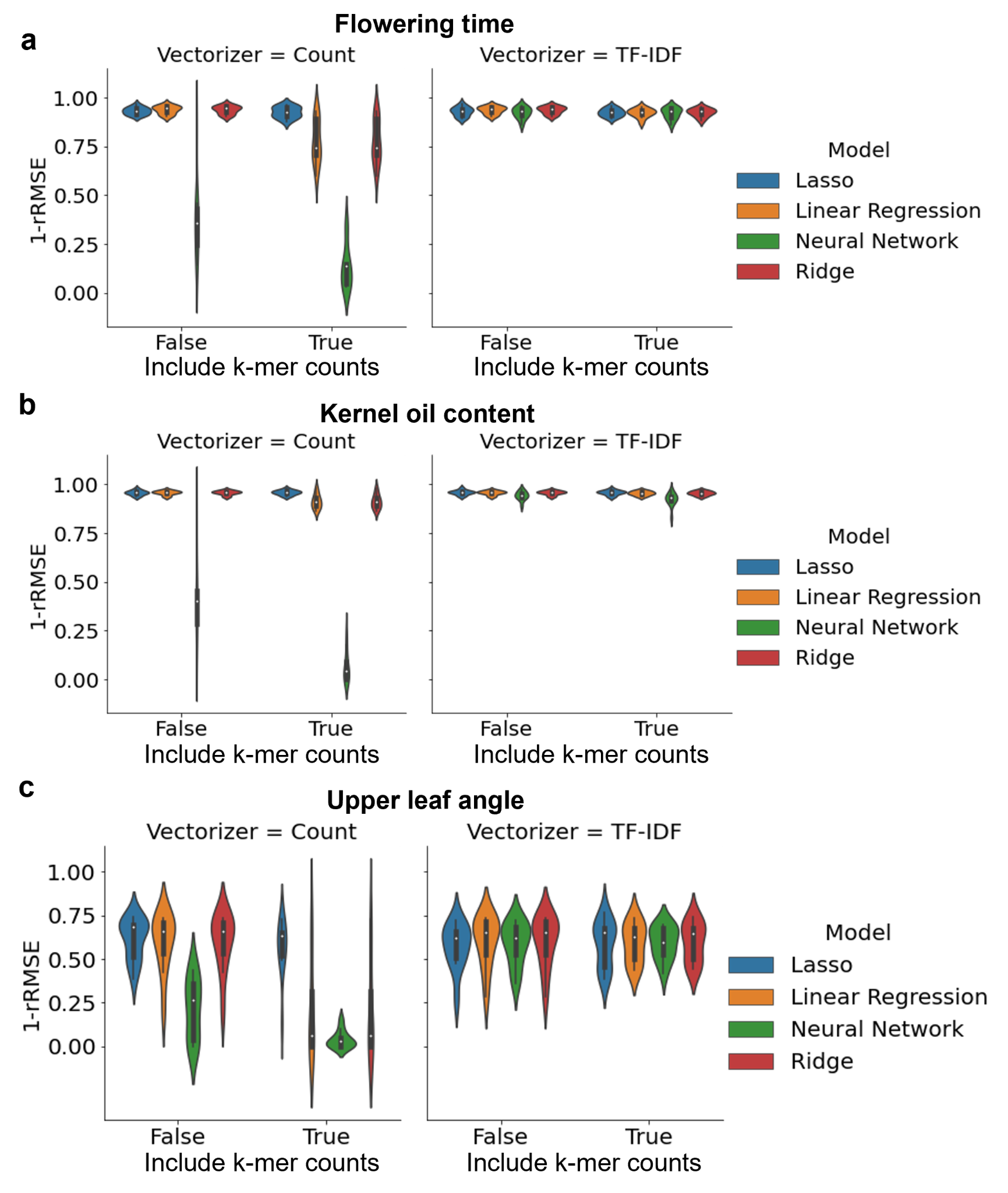

Supplementary Fig. 10. Performance of different machine learning model variations. (a) Flowering time, (b) kernel oil, and (c) upper leaf angle.TF-IDF = term frequency-inverse document frequency. 1-rRMSE = one minus relative root means squared error.
